# Branch-specific axon pruning induced by Dpr4/DIP-θ transneuronal interactions

**DOI:** 10.64898/2026.03.29.715068

**Authors:** Hagar Meltzer, Shirel Shahar, Alina P. Sergeeva, Bavat Bornstein, Gal Shapira, Phinikoula S. Katsamba, Seetha Mannepalli, Fabiana Bahna, Noa Moreno, Idan Alyagor, Voctoria Berkun, Timothy Currier, Lawrence Shapiro, Barry Honig, Oren Schuldiner

**Author notes:** These authors contributed equally (Hagar Meltzer, Shirel Shahar).

## Abstract

Neuronal remodeling is a conserved, late developmental mechanism to refine neural circuits. Although remodeling typically occurs with remarkable spatiotemporal precision, its underlying molecular mechanisms remain poorly understood. In the *Drosophila* mushroom body (MB) circuit, γ-Kenyon cells (γ-KCs) undergo stereotyped remodeling during metamorphosis, in which they prune their larval vertical and medial axonal branches and subsequently regrow a medial, adult-specific branch. Our previous transcriptional profiling of developing γ-KCs revealed dynamic expression of Defective proboscis extension response (Dpr) proteins and their binding partners, Dpr-interacting proteins (DIPs), members of the Immunoglobulin (Ig) superfamily. Despite their established roles in neurodevelopment, how Dpr/DIPs function - given their lack of intracellular domains - remains unclear. Here, we show that overexpression of Dpr4 in developing γ-KCs cell-autonomously inhibits axon pruning. Strikingly, this effect is branch-specific: the vertical axonal branch fails to prune, while the medial branch prunes normally. To our knowledge, this represents the first demonstration of branch-specific control of pruning in this system. Moreover, the adult medial branch regrows normally, indicating that pruning and regrowth are independently regulated at the level of individual branches. We demonstrate that this unique branch-specificity arises from trans-neuronal interactions between Dpr4 in γ-KCs and DIP-θ in dopaminergic neurons that selectively innervate the vertical larval MB lobe. Furthermore, our findings suggest that this phenotype relies on an Ig2 domain of a Dpr family member, implying the involvement of a third binding partner. Leveraging this robust overexpression phenotype to probe downstream mechanisms, we find that loss of the transmembrane adhesion protein N-Cadherin suppresses the Dpr4-induced pruning defect. Together, our findings highlight the local impact of Dpr/DIP-mediated trans-neuronal interactions on the spatial regulation of remodeling, and provide genetic evidence implicating N-Cadherin as a potential downstream mediator of Dpr/DIP function within a developing neural circuit.

## Introduction

Shaping the complex yet precise connectivity of the mature nervous system relies on a combination of degenerative and regenerative developmental events. Collectively referred to as neuronal remodeling, such processes represent an evolutionary conserved strategy to refine neural circuits, and often include elimination of exuberant connections followed by regrowth to adult-specific targets. In humans, massive elimination of embryonic connections occurs during the first two years of life, and continues, to a lesser extent, until after adolescence (Luo & O’Leary, 2005; Schuldiner & Yaron, 2015; Riccomagno & Kolodkin, 2015). Defects in remodeling have been associated with various neurodevelopmental conditions, including schizophrenia and autism spectrum disorder (Sekar et al., 2016; Thomas et al., 2016; Cocchi et al., 2016). Stereotypic neuronal remodeling follows a fixed course in space and time, thus allowing us to predict exactly which neural processes will be eliminated, to what extent, and at which developmental stage. Despite the widespread occurrence and fundamental nature of stereotypic remodeling, our understanding of its spatial regulation, and specifically why predetermined branches, or branch parts, are pruned while others are spared, remains incomplete (Furusawa & Emoto, 2020, 2021; Meltzer & Schuldiner, 2022).

During the metamorphosis of *Drosophila melanogaster*, its entire nervous system undergoes stereotypic rearrangements (Truman, 1990; Yaniv & Schuldiner, 2016). Combined with its unparalleled neurogenetic toolbox, *Drosophila* is an ideal model to explore the molecular mechanisms governing neuronal remodeling. We focus on the mushroom body (MB), a complex circuit in the fly brain that functions as a center of associative learning and memory. The MB is comprised of 3 types of sequentially-born intrinsic neurons termed Kenyon cells (KCs), out of which only the first-born γ-KCs undergo extensive axonal remodeling during metamorphosis. In the embryonic and early larval stages, γ-KCs project axons that bifurcate to form vertical and medial branches. However, in the early pupal stage - approximately 6 hours after puparium formation (h APF) - both axonal branches begin to prune, and complete their elimination up to their branchpoint by 18h APF. At 24h APF, γ-KCs initiate regrowth of an adult-specific, medially-projecting axonal branch (Fig. 1A; Lee et al., 1999; Yu & Schuldiner, 2014; Yaniv & Schuldiner, 2016; Lin, 2023). The stereotypic pruning and regrowth of MB axons captures the spatiotemporal precision that often characterizes remodeling processes, but the underlying mechanisms remain unclear.

**Figure 1.**
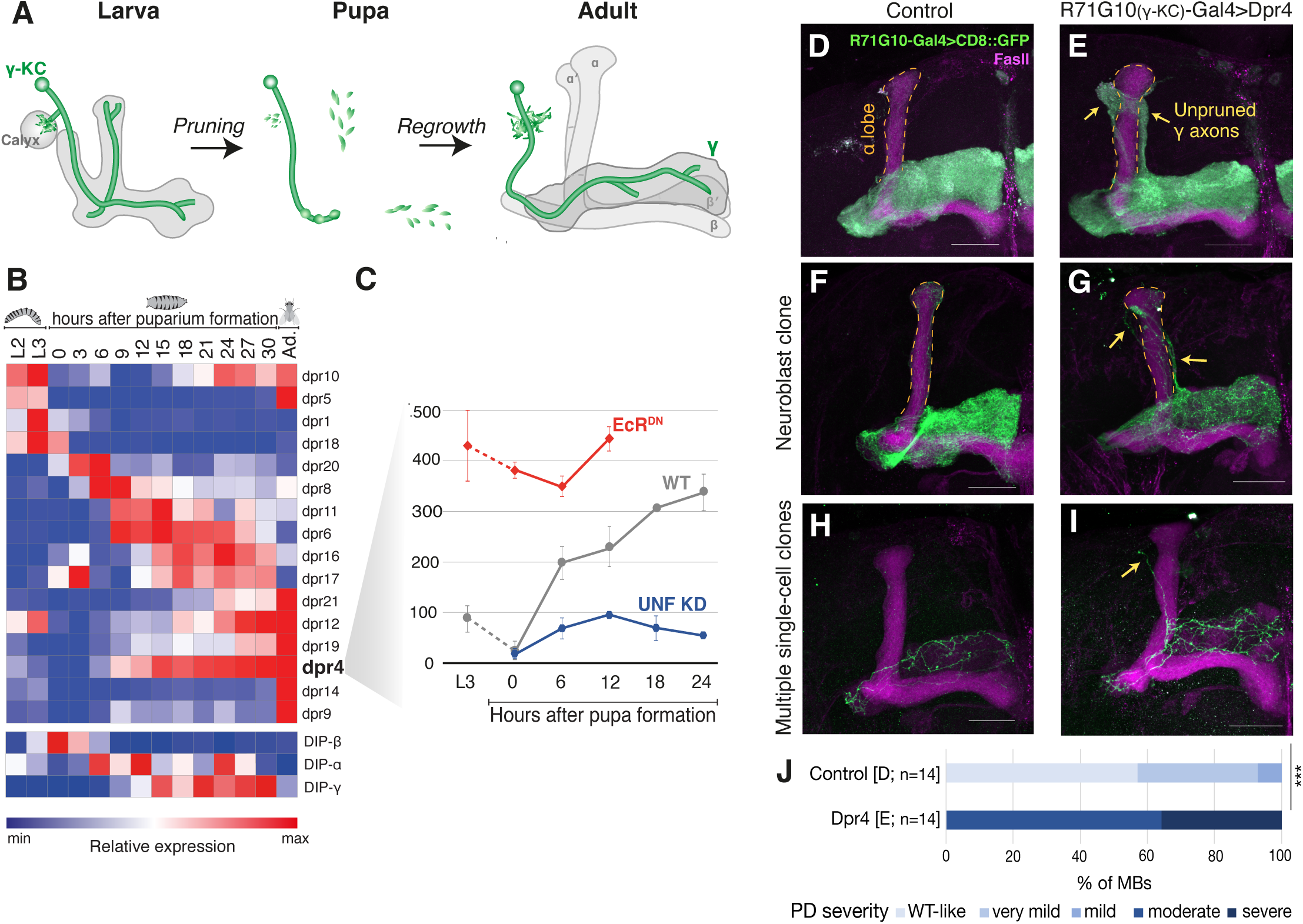
Overexpression of Dpr4 cell-autonomously inhibits axon pruning of mushroom body γ-KCs. (A) Scheme of MB remodeling. When they initially grow, γ-KCs (green) project axons that bifurcate to form vertical and medial branches. During early pupal stages, dendrites (in the calyx) are pruned completely, and axonal branches are pruned up to their branchpoint. Later, an adult-specific medial axon regrows. h APF; hours after puparium formation. (B) Heatmap depicting the transcriptional profiles of Dprs and DIPs along γ-KC development (Alyagor et al., 2018; Bornstein et al., 2021). Red and blue represent high and low relative expression, respectively. (C) Expression plot of Dpr4 in developing γ-KCs, either WT (grey), upon expression of a dominant negative variant of Ecdysone receptor (EcR^DN^; red), or upon knockdown of unfulfilled (UNF KD; blue). The Y-axis represents gene expression levels in arbitrary units. (D-E) Confocal z-projections of adult MBs in which R71G10-Gal4 (expressed in γ-KCs and stochastically also in α/β-KCs) drives expression of membranal GFP (CD8::GFP; green; D is control), or additionally driving overexpression of Dpr4 (E). (F-I) Confocal z-projections of adult MBs with neuroblast (F-G) or multiple single-cell (H-I) MARCM clones of γ-KCs, positively labeled by R71G10-Gal4-driven membranal GFP (CD8::GFP; green; F,H is control) or additionally overexpressing Dpr4 (G,I). Magenta is FasII staining, which strongly labels α/β-axons and weakly labels γ-axons. The α lobe is outlined in orange. Yellow arrows point to unpruned larval axons. Scale bar represents 30μm. (J) Quantification of the pruning defect severity in D-E, ranging from WT-like morphology (score=0; light blue) to severe pruning defect (score=4; dark blue). The percentage of MBs displaying each score is represented. Mann-Whitney U-test \*\*\**p* = 3.5 ⨉ 10^-6^.

Spatial regulation of remodeling could be achieved by cell intrinsic mechanisms and/or via extrinsic cues from adjacent cells or the extracellular matrix (Meltzer & Schuldiner, 2020). Cell surface proteins (CSPs), which are potential mediators of such cell-cell interactions, are dynamically expressed in developing γ-KCs, as evident in the transcriptional atlas we previously generated (Alyagor et al., 2018). Specifically, many members of the Defective proboscis extension response (Dpr) family display highly dynamic expression patterns during the remodeling timeframe (Fig. 1B). This family consists of 21 immunoglobulin superfamily (IgSF) proteins that form an elaborate interactome, known as the ‘Dprome’, with 11 Dpr-interacting proteins (DIPs; Cosmanescu et al., 2018; Ozkan et al., 2013; Carrillo et al., 2015). Over the past decade, Dpr/DIP interactions were shown to mediate various processes of target recognition and synaptic wiring in developing *Drosophila* circuits (e.g., Barish et al., 2018; Xu et al., 2018; Ashley et al., 2019; Venkatasubramanian et al., 2019; Xu et al., 2022; Morano et al., 2025). In the MB, we previously established a role for trans-neuronal Dpr/DIP interactions in circuit reassembly during γ-axon regrowth (Bornstein et al., 2021). However, the molecular mechanisms and signaling pathways by which Dpr/DIPs exert their function in the MB, as well as in other developing circuits, remain unknown. As recent evidence suggests that most, if not all, Dprome members are Glycosylphosphatidylinositol (GPI)-anchored proteins (Lobb-Rabe et al., 2024), intracellular signaling is likely mediated by additional molecules, which are yet to be identified.

Here, we focus on Dpr4, which is downregulated before the onset of γ-KC axon pruning. We demonstrate that its overexpression in γ-KCs results in a unique phenotype in which pruning of the vertical, but not medial, axonal branches is inhibited, without affecting regrowth of the adult medial branch. We show that this spatial specificity is due to ectopic trans-neuronal interactions with vertical lobe-innervating dopaminergic neurons that endogenously express its binding partner DIP-θ. Finally, we provide genetic evidence of the adhesion molecule N-Cadherin as a potential functional mediator of this branch-specific pruning defect. Our findings shed new light on the function of Dpr/DIPs in the spatial regulation of neuronal remodeling, and suggest a novel effector for their neurodevelopmental function.

## Results

### Overexpression of Dpr4 cell-autonomously inhibits axon pruning of mushroom body γ-KCs

Our previous transcriptional profiling of developing MB γ-KCs (Alyagor et al., 2018; Bornstein et al., 2021) revealed interesting expression dynamics of many Dprome members (Fig. 1B). Among them is Dpr4, which is low at the onset of metamorphosis (0h APF), but gradually increases its expression upon the initiation of γ-axon pruning (at 6h APF) and remains high during axon regrowth (initiated at 24h APF; Fig. 1C). Additionally, the expression of Dpr4 is affected by perturbation of key regulators of axon pruning (Ecdysone receptor; EcR; Lee et al., 2000), or regrowth (unfulfilled; UNF; Yaniv et al., 2012): expression of a dominant negative variant of EcR EcR^DN^) results in elevated levels of Dpr4 throughout development, while knockdown of UNF dramatically downregulates Dpr4 expression (Fig. 1C). These combined expression profiles, which are similar to those of Dpr12 (Bornstein et al., 2021), made us hypothesize that Dpr4 is specifically silenced before axon pruning, or required for axon regrowth, or both. We therefore knocked down Dpr4 within γ-KCs using the γ-specific driver R71G10-Gal4, but observed no effect on axon pruning or regrowth (SFig. 1). Importantly, this result does not rule out the involvement of Dpr4 in γ-axon regrowth, as the extensive redundancies within the Dprome may require simultaneous knockdown of multiple Dprs to achieve phenotypic effects. We next utilized the FlyORF collection (Bischof et al., 2013) to overexpress Dpr4 in γ-KCs throughout development using R71G10-Gal4. Indeed, this resulted in a severe γ-axon pruning defect, manifested as persistence of vertically-projecting γ-axons (SFig. 2A-B). However, when we stained for the HA tag inserted at the C-terminus of the FlyORF transgene, we could only detect the HA signal in the γ cell bodies and not in the axons (SFig. 2B’). This is in line with Dprs being GPI-anchored proteins (Lobb-Rabe et al., 2024), and therefore a C-terminal tag is expected to be cleaved before the protein reaches the plasma membrane. We therefore generated a new tagged Dpr4 transgene (UAS-HA.Dpr4) in which HA is inserted at the N-terminus, downstream of the predicted signal peptide and upstream of the first Ig domain (SFig. 2E). Overexpression of this transgene using R71G10-Gal4 recapitulated the pruning defect (Fig. 1D-E, quantified in J), and the HA signal was observed throughout the entire neuron, including the axonal lobes, in both larval and adult MBs (SFig. 1C-D). Notably, this effect seems to be specific to Dpr4, as γ-KCs overexpressing other Dprs, including Dpr10 and Dpr12, prune normally (SFig. 3).

To explore the nature of the pruning defect, we overexpressed Dpr4 in clones using the MARCM (Mosaic analyses with a repressible cell marker; Lee & Luo, 1999) technique. Positively labelled neuroblast clones - in which only a fourth of the γ-KC population overexpresses Dpr4 – exhibited axon pruning defects (Fig. 1F-G). While we were unable to observe single cell clones displaying a pruning defect, in a few MBs with multiple single-cell clones we could detect one cell retaining an unpruned larval vertical axon (Fig. 1H-I). These results indicate that Dpr4 overexpression inhibits γ-axon pruning in a cell-autonomous manner.

### Dpr4 inhibits axon pruning in a branch-specific manner, while regrowth remains unaffected

Developmental analysis revealed that at 24h APF, the timepoint by which the larval axonal lobes have normally completed pruning up to their branchpoint and are initiating regrowth, Dpr4-overexpressing γ-KCs displayed normal pruning of the medial lobe, but a virtually intact vertical lobe (Fig. 2A,C; quantified in SFig. 4). This lobe-specific pruning defect is unusual, as in all the cases we encountered in the past, pruning was similarly disrupted in both lobes. This can be seen, for example, in the pruning defect caused by overexpression of EcR^DN^ in γ-KCs (Fig. 2B; quantified in SFig. 4). We conclude that Dpr4 inhibits pruning of γ-axons in a unique, branch-specific manner. To the best of our knowledge, this is the first description of such a phenotype in this system.

**Figure 2.**
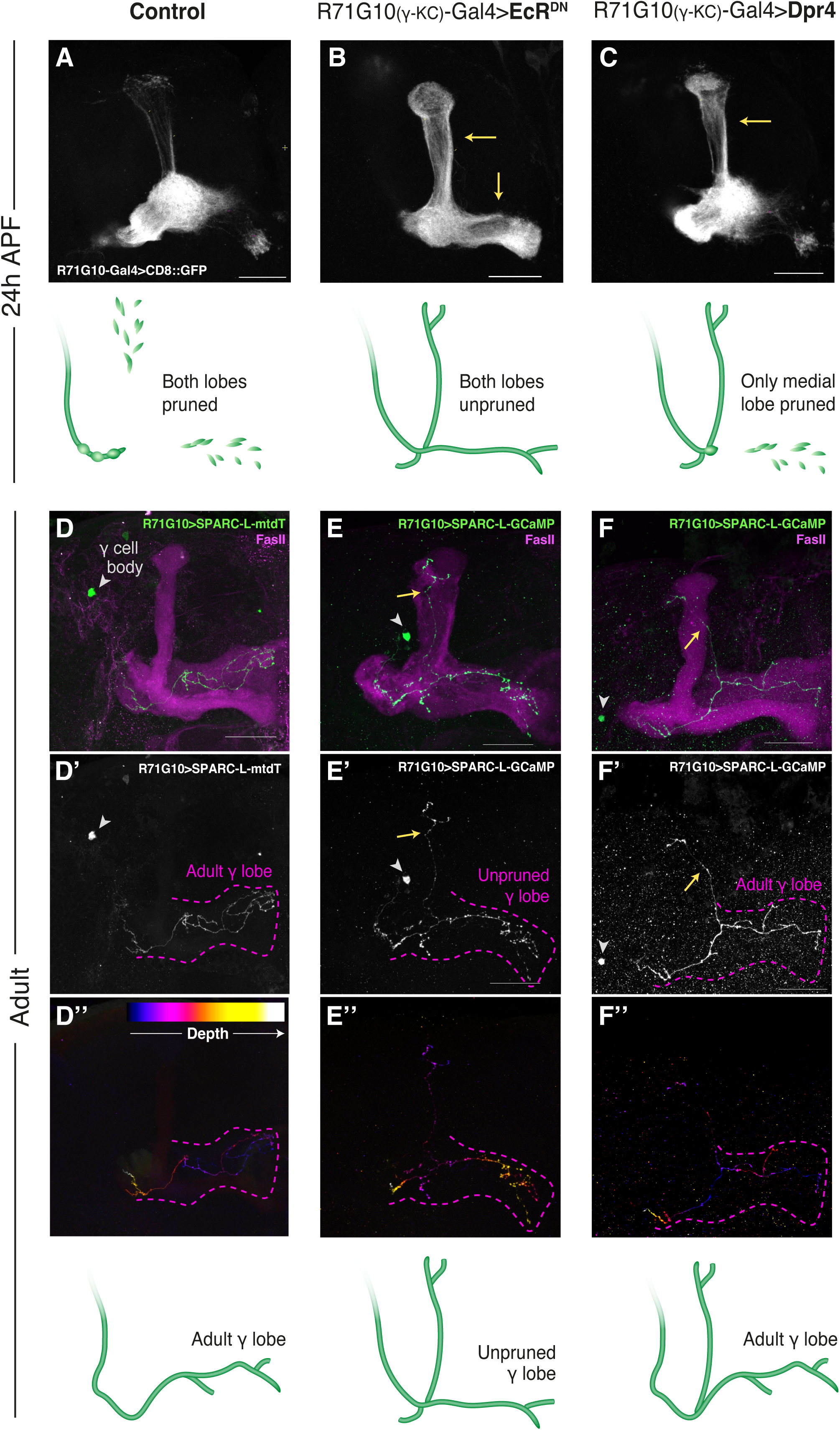
Dpr4 inhibits axon pruning in a branch-specific manner, while regrowth remains unaffected. (A-C) Confocal z-projections of MBs at 24 hours after puparium formation (h APF), in which R71G10-Gal4 drives expression of membranal GFP (CD8::GFP; greyscale) in γ-KCs, in control (A) or also driving overexpression of either a dominant negative variant of EcR (EcR^DN^; B) or Dpr4 (C). Yellow arrows point to unpruned γ lobes. Scale bar represents 30μm. The illustrations below represent the observed phenotypes. (D-F) Confocal z-projections of adult MBs in which R71G10-Gal4 drives expression of myr::tdTomato (mtdT; stained with anti-RFP antibody; green in D, greyscale in D’, color-coded based on z-depth within a sub-stack of the axonal lobes in D’’), or GCaMP8m (stained with anti-GFP antibody; green in E-F, greyscale in E’-F’, color-coded based on z-depth within a sub-stack of the axonal lobes in E’’-F’’) in a single γ-KC, using the SPARC-L strategy (Currier et al., 2025). Additionally, R71G10-Gal4 drives expression of EcR^DN^ (E) or Dpr4 (F) in the entire γ-KC population (not only in the marked cell). Magenta is FasII staining, which strongly labels α/β-axons and weakly labels γ-axons. Yellow arrows point to single unpruned γ vertical axons. Arrowheads point to the γ-cell bodies. The medial γ lobe is outlined in magenta in E’-G’ and E’’-G’’. Scale bar represents 30μm. The illustrations below represent the observed phenotypes.

Pruning of the vertical and medial axonal branches is followed by regrowth of an adult-specific axonal branch that only projects medially. Importantly, the adult medial branch is anatomically distinct from the larval one, and occupies a more dorsal and anterior location in the MB. The phenotype of Dpr4 overexpression, in which only the medial branch prunes, provides us with the unique opportunity to test whether regrowth of the medial branch can occur despite persistence of the vertical larval branch. We therefore used an ultra-sparse variant of the SPARC technique, ‘layered’ SPARC (SPARC-L; Currier & Clandinin, 2025) that allows extremely sparse labeling of neurons at up to single cell resolution (while an independent transgene of interest can be simultaneously expressed in the entire Gal4 domain). WT single γ-KCs show the typical location of the adult-specific medial branch (Fig. 2D-D’’), while in EcR^DN^-expressing single γ-KCs the medial branch occupies the larval location, which is spatially distinct and occupies a deeper and more ventral position compared to the adult branch (Fig. 2E-E’’). Remarkably, upon Dpr4 overexpression, we were able to identify single γ-KCs in which the vertical axonal branch retained its larval state, while the medial branch occupied its adult-specific location (Fig. 2F-F’’). These tempo-chimeric neurons, of both larval and adult features, suggest that axon pruning and regrowth are spatially separable processes that can be independently controlled at the level of a single branch.

### Dpr4 inhibits axon pruning by binding to one (or more) of its cognate DIPs

We hypothesized that the branch-specific pruning defect induced by Dpr4 overexpression arises from *trans* interactions with one or more of its cognate DIPs in interacting neurons. *In vitro* assays found that Dpr4 can bind DIP-θ and DIP-η (Carrillo et al., 2015) as well as DIP-ι (Cosmanescu et al., 2018). To disrupt these interactions, we designed three single point mutations (I87D, Y95D and Q130D) located in the first immunoglobulin domain (Ig1) of Dpr4 (Fig. 3A). These amino acid substitutions replace buried hydrophobic (Ile, Tyr) or polar (Gln) residues at the binding interface with negatively charged aspartates, thereby introducing a desolvation penalty predicted to weaken Dpr4::DIP-θ/η/ι binding. Surface plasmon resonance (SPR) binding assays confirmed that the Dpr4 mutants I87D, Y95D and Q130D abrogated binding to the cognate Dpr4 partners, DIP-η and DIP-θ, without enabling interactions with the non-cognate partner DIP-α (SFig. 5 and Table 1). Following these *in vitro* validations, we tested two of the mutants *in vivo* within the developing MB. We generated transgenic flies expressing HA-tagged Dpr4 variants carrying either the I87D or the Y95D mutation. Overexpression of these transgenes in γ-KCs resulted in widespread localization of the mutated protein throughout the neuron (as evident by the HA tag; Fig. 3D’-E’). However, unlike overexpression of WT Dpr4, which consistently caused prominent pruning defects (Fig. 3C), overexpressing the I87D or Y95D mutant transgenes did not produce any pruning abnormalities (Fig. 3D-E, quantified in F). These findings support our hypothesis that pruning inhibition depends on Dpr4 interactions with one or more of its cognate DIPs.

**Figure 3.**
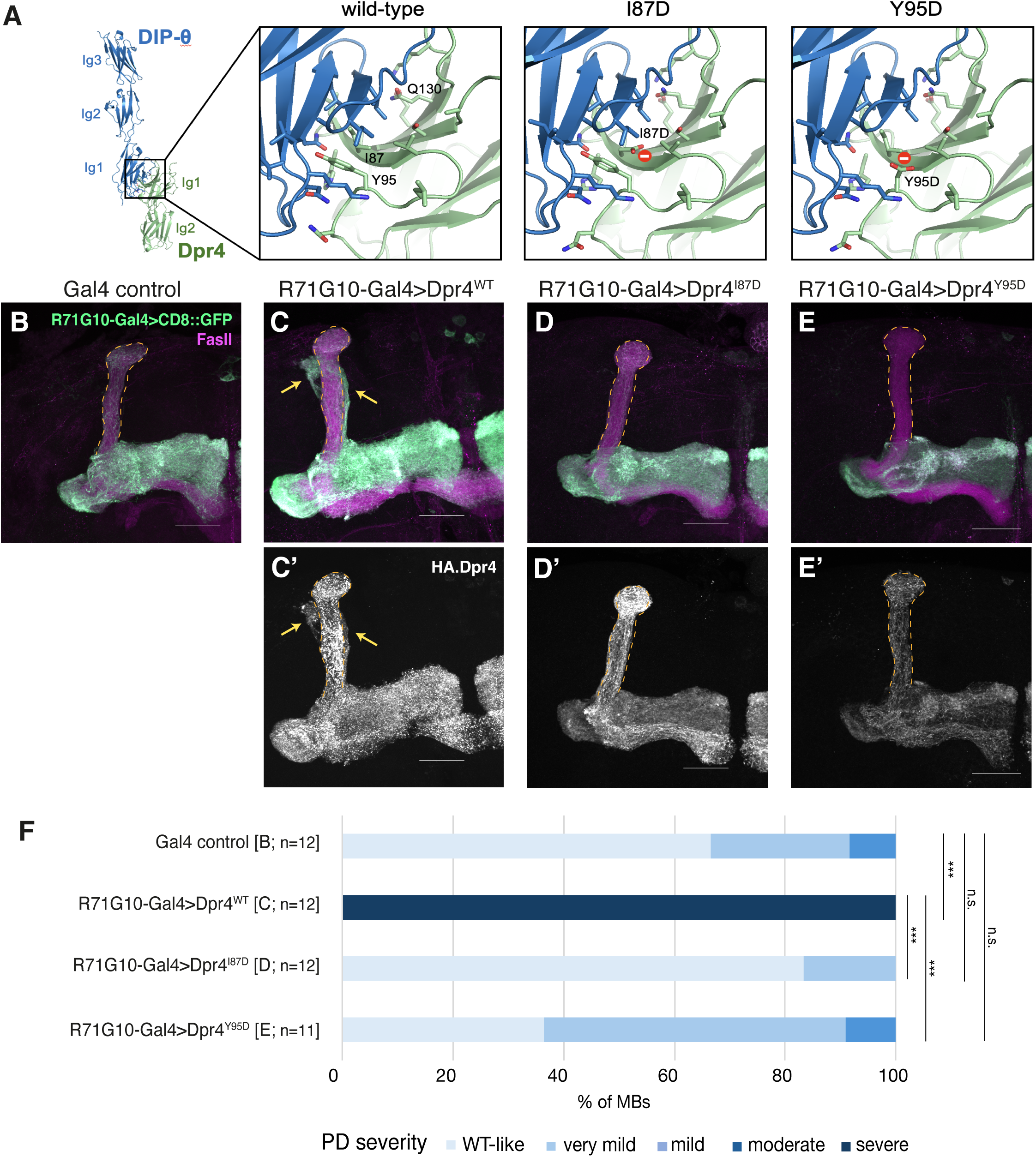
Dpr4 inhibits pruning by binding one (or more) of its cognate DIPs. (A) Ribbon representation model of the Dpr4::DIP-θ complex, with insets highlighting interfacial residues (as sticks) in WT versus I87D/Y95D mutants, showing mutation-induced buried unsatisfied charges (as red spheres). (B-E) Confocal z-projections of adult MBs in which R71G10-Gal4 drives expression of membranal GFP in γ-KCs (CD8::GFP; green), either control (B) or also overexpressing WT Dpr4 (C) or Dpr4 harboring the I87D (D) or Y95D (E) mutations. Magenta is FasII staining, which strongly labels α/β-axons and weakly labels γ-axons. Staining for the HA tag inserted in the N-terminus of the WT/mutant Dpr4 protein is shown in greyscale in C’-E’. Note that R71G10-driven HA.Dpr4 (and to a lesser extent CD8::GFP) is also expressed in the α lobe which is outlined in orange. Yellow arrows indicate unpruned γ-axons. Scale bar represents 30μm. (F) Quantification of the pruning defect severity in B-E, ranging from WT-like morphology (score=0; light blue) to severe pruning defect (score=4; dark blue). The percentage of MBs displaying each score is represented. Kruskal-Wallis test *p* = 3.724 ⨉ 10^-7^; Mann-Whitney U-test: [C] vs. [D] \*\*\**p* = 1.9 ⨉ 10^-5^; [C] vs. [E] \*\*\**p* = 2.1 ⨉ 10^-5^; [B] vs. [C] \*\*\**p* = 1.9 ⨉ 10^-5^; [B] vs. [D] not significant (n.s.) *p*=0.339; [B] vs. [E] not significant (n.s.) *p*=0.258.

**Table 1.** Equilibrium-binding K_D_s of WT/mutant Dpr4 interactions with cognate (DIP-ρι/8) and non-cognate (DIP-α) DIPs. See also SFig. 5 for sensorgrams and binding isotherms.

|  | **K<sub>D</sub> (μM)** |  |  |
| --- | --- | --- | --- |
| | DIP-η | DIP-θ | DIP-α |
| Dpr 4 WT | 105(1) | 46.7(4) | >500 |
| Dpr 4 I87D | >500 | >500 | >500 |
| Dpr4 Y95D | >500 | >500 | >500 |
| Dpr4 Q130D | >500 | >500 | >500 |

### Pruning of the vertical lobe is inhibited by trans-neuronal interactions between Dpr4 in γ-KCs and DIP-θ in dopaminergic neurons

The spatially-restricted pruning defect, despite the widespread presence of the transgenic Dpr4 protein in both the vertical and medial larval lobes (SFig. 2D’), suggested that the interacting DIP is expressed in neurons that specifically innervate the vertical γ-lobe. In addition to the intrinsic KCs, the MB circuit consists of extrinsic neurons, including MB output neurons (MBONs), which relay the sensory information from KCs to other brain regions, and two distinct clusters of dopaminergic neurons (DANs), which modulate the KC-MBON synapses (Tanaka et al., 2008; Aso et al., 2014; Saumweber et al., 2018; Truman et al., 2023). In both larval and adult MBs, the vertical lobes are innervated by DANs of the PPL1 cluster (also known as DL1 in larvae; Weber et al., 2025), while modulatory innervation of the medial lobes is via the PAM cluster (or pPAMs in larvae; Fig. 4A). We therefore hypothesized that overexpressed Dpr4 in γ-KCs forms ectopic interactions with one of its cognate DIPs within PPL1-DANs, thereby inhibiting the normal progression of pruning only within the vertical lobe. We examined an existing transcriptional dataset of adult MB neurons (Aso et al., 2019) and found that out of Dpr4’s cognate DIPs, DIP-θ is the only one consistently expressed within PPL1-DANs. While we cannot necessarily deduce about the developmental expression from the adult profile, this motivated us to focus on DIP-θ. We therefore used a DIP-θ-T2A-Gal4 line, expected to express Gal4 in the cells that endogenously express DIP-θ, to drive membranal GFP. At 8h APF, shortly after the initiation of pruning, we observed numerous GFP-positive processes surrounding the vertical axonal lobe. Furthermore, we could clearly detect GFP signal in the cell bodies of 4 neurons of the PPL1 cluster, which was co-localized with the dopaminergic marker tyrosine hydroxylase (TH; SFig. 6). Our findings suggest that DIP-θ is expressed in specific PPL1-DANs innervating the vertical lobe within the pruning timeframe.

**Figure 4.**
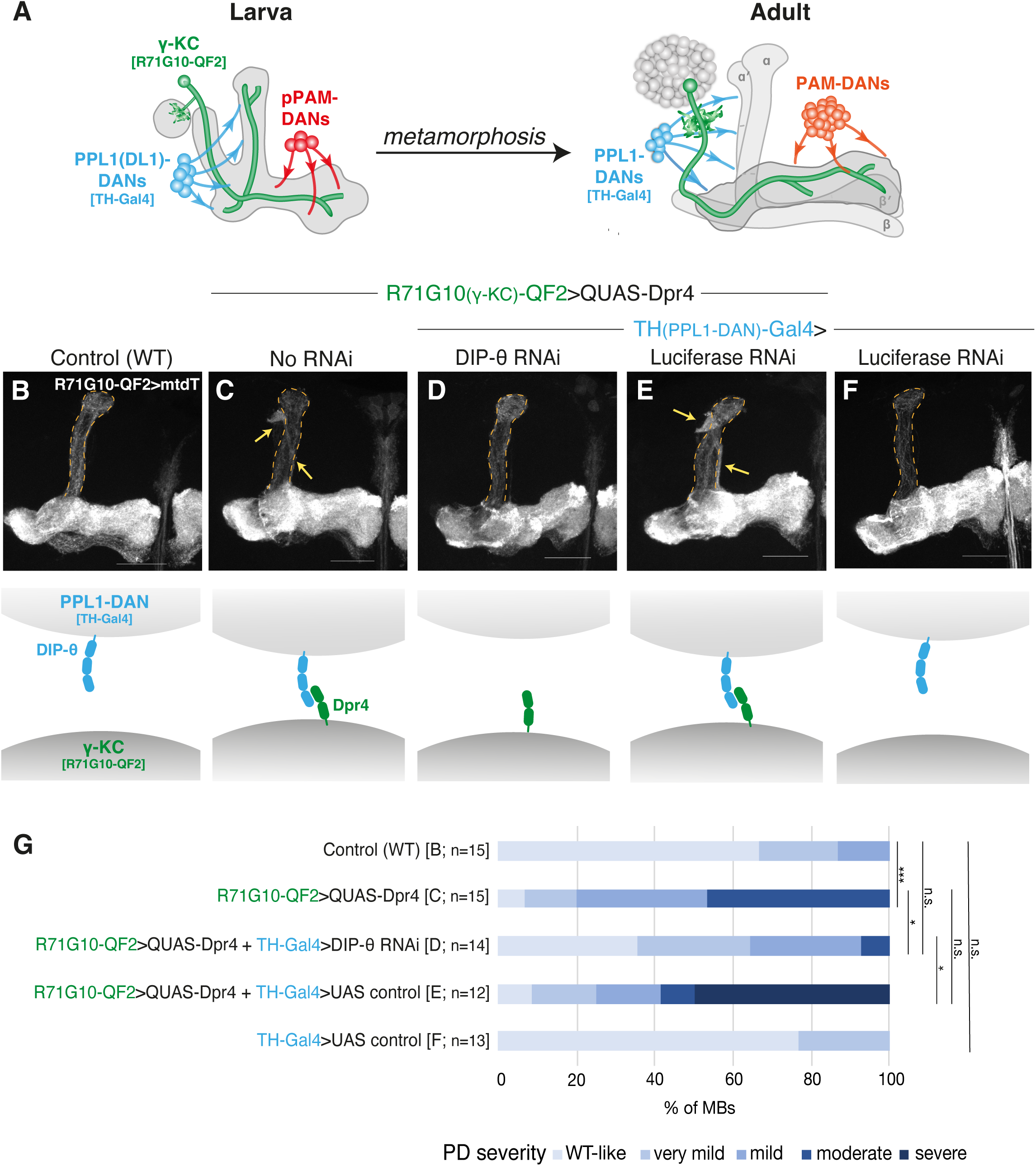
Pruning of the vertical lobe is inhibited by trans-neuronal interactions between Dpr4 in γ-KCs and DIP-θ in dopaminergic neurons. (A) Scheme of MB remodeling, including dopaminergic neurons (DANs) of the PPL1 (blue) and PAM (red) clusters, which largely innervate the vertical and medial lobes, respectively, in both larval and adult MBs. (B-F) Confocal z-projections of adult MBs in which R71G10-QF2 drives expression of myr::mtdTomato (mtdT; greyscale) in γ-KCs, and to a lesser extent in α/β-KCs. (B) is control while in (C-E) R71G10-QF2 also drives Dpr4 overexpression. In (D-F) the PPL1-DAN-specific TH-Gal4 drives expression of RNAi targeting DIP-θ (D), or of a control UAS trasngene (UAS-luciferase; E-F). The α lobe is outlined in orange. Yellow arrows indicate unpruned γ-axons. Scale bar represents 30μm. The schemes below illustrate presumed protein expression and interactions on γ-KC and PPL1-DAN membranes in each condition. (G) Quantification of the pruning defect severity in B-F, ranging from WT-like morphology (score=0; light blue) to severe pruning defect (score=4; dark blue). The percentage of MBs displaying each score is represented. Kruskal-Wallis test p<0.001; Mann-Whitney U-test: [B] vs. [C] \*\*\**p*<0.001; [C] vs. [D] \**p*=0.011; [D] vs. [E] * *p*=0.011; [B] vs. [D] not significant (n.s.) *p*=0.103; [C] vs. [E] not significant (n.s.) *p*=0.203; [B] vs. [F] not significant (n.s.) *p*=0.467.

If pruning is indeed inhibited due to ectopic interactions between Dpr4 in γ-KCs and DIP-θ in PPL1-DANs, then downregulation of DIP-θ in PPL1-DANs is expected to suppress the Dpr4-induced pruning defect. Experimental validation of our hypothesis requires simultaneous manipulation of two distinct cell populations. We thus generated a QUAS-Dpr4 transgene and confirmed that, like its UAS counterpart, it inhibits pruning when overexpressed using the γ-KC driver R71G10-QF2 (Fig. 4B-C, quantified in G). When, on this background, we independently used the Gal4/UAS system to specifically knockdown DIP-θ in PPL1-DANs (using the TH-Gal4 driver), this was sufficient to significantly suppress the Dpr4-induced pruning defect (Fig. 4D, quantified in G). As expected, using TH-Gal4 to drive a UAS-Luciferase construct (which serves as control for pVALIUM10 TRIP lines) had no suppressive effect and the Dpr4-induced pruning defect persisted (Fig. 4E; we also confirmed that this construct itself does not inhibit pruning, Fig. 4F, quantified in G). Together, our experiments indicate that ectopic trans-neuronal interactions between overexpressed Dpr4 in γ-KCs, and endogenously-expressed DIP-θ in PPL1-DANs, inhibit axon pruning in a spatially restricted manner.

### The Dpr4-induced pruning defect depends on an Ig2 domain of a Dpr family member

Dprs are known to bind DIPs exclusively via the Ig1 domains of both proteins (Cosmanescu et al., 2018; Sergeeva et al., 2020). Indeed, specifically mutating Dpr4 in its Ig1 interface with DIP-θ abolished its inhibitory effect on pruning (Fig. 3). However, the functional significance of the Ig2 domain, present in all Dprome members, remains largely unknown. To explore the potential role of Ig2 in mediating the Dpr4-induced pruning defect, we designed and generated chimeric Dpr4 transgenes in which its Ig2 is replaced with the Ig2 of Dpr12 or with the Ig2 of the IgSF adhesion molecule Fasciclin II (FasII). When overexpressed in γ-KCs, the Dpr12 Ig2 replacement transgene consistently showed prominent pruning defects - identical in severity to those of WT Dpr4 overexpression (Fig. 5A vs. B, quantified in D). However, replacement with the FasII Ig2 displayed pruning defects that were significantly weaker than those observed in the WT or Dpr12-replaced Ig2 (Fig. 5C, quantified in D). Immunostaining of non-permeabilized brains confirmed that the overexpressed chimeric Dpr4 proteins are localized to the cell surface of γ-KCs, similarly to the WT overexpression transgene (SFig. 7). Taken together, our results suggest that the strong inhibition of pruning by Dpr4 relies on an Ig2 domain which seems to be conserved among Dprs but not among other IgSF adhesion molecules.

**Figure 5.**
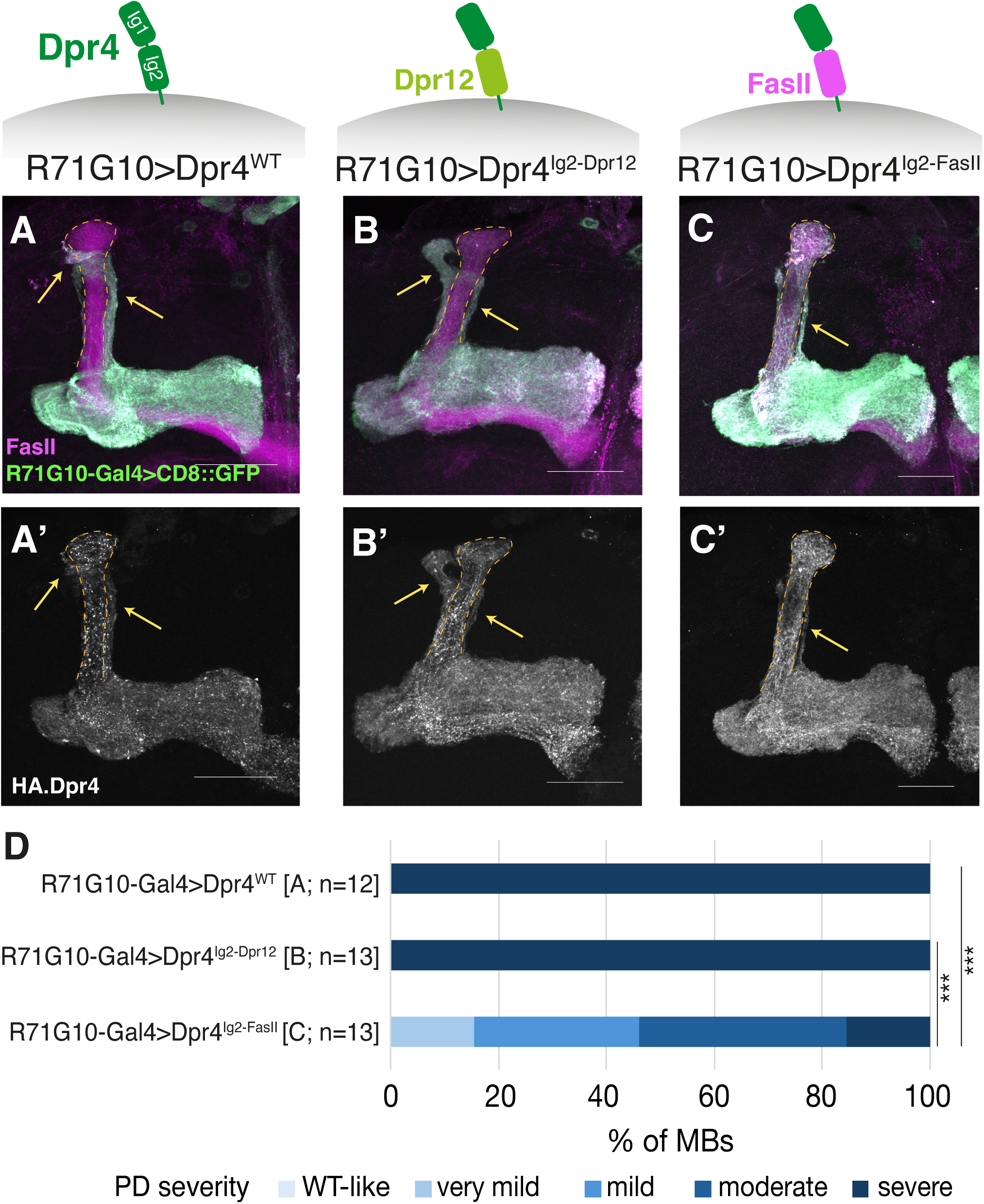
The Dpr4-induced pruning defect depends on an Ig2 domain of a Dpr family member. (A-C) Confocal z-projections of adult MBs in which the R71G10-Gal4 drives expression of membranal GFP (CD8::GFP; green) as well as WT Dpr4 (A), or Dpr4 in which the entire Ig2 is replaced with the Ig2 of Dpr12 isoform C (B) or of FasII isoform C (C). Magenta is FasII staining, which strongly labels α/β-axons and weakly labels γ-axons. Staining for the HA tag inserted in the N-terminus of the WT/mutant Dpr4 protein is shown in greyscale in A’-C’. Note that R71G10-driven HA.Dpr4 (and to a lesser extent CD8::GFP) is also expressed in the α lobe which is outlined in orange. Yellow arrows indicate unpruned γ-axons. Scale bar represents 30μm. (D) Quantification of the pruning defect severity in A-C, ranging from WT-like morphology (score=0; light blue) to severe pruning defect (score=4; dark blue). The percentage of MBs displaying each score is represented. The same 12 Dpr4^WT^ images were also used for quantification in Figure 3 (as the genotype of Fig. 3C and Fig. 5A is identical). Kruskal-Wallis test *p* = 8.113 ⨉ 10^-7^; Mann-Whitney U-test: [B] vs. [C] \*\*\**p* = 7.8 ⨉ 10^-5^; [A] vs. [C] *** *p* = 7.8 ⨉ 10^-5^; [A] and [B] share an identical (most severe) score so a direct comparison between them was not applicable.

### Loss of N-Cadherin suppresses the Dpr4-induced pruning defect

Dprs and DIPs were shown to be GPI-anchored proteins (Cheng et al., 2019; Lobb-Rabe et al., 2024), and none encode for a sufficiently long intracellular domain. Therefore, it is highly likely that Dpr/DIP signaling occurs via yet unknown interacting proteins. The importance of the Ig2 domain in the *C. elegans* homolog of DIPs - RIG-5 - was recently highlighted in an extracellular interactome study, which found it likely mediates binding to the nematode homologs of *Drosophila* Nrx-IV and Lar (Nawrocka et al., 2026). Importantly, the same study also demonstrated *in vitro* binding of RIG-5 to HMR-1, which is the nematode homolog of *Drosophila* N-Cadherin (CadN), although their interaction interface was not specified. The functional importance of Dpr4 Ig2 (Fig. 5) makes Dpr4 overexpression an exceptional platform to identify interacting proteins, given its robust and consistent phenotype, which can be used as a baseline for suppression assays. Out of the candidates in the Nawrocka et al. study, our attention was immediately drawn to CadN, as it resonated our recently published observation that it is required for γ-axon regrowth (Fahdan et al., 2026) - which coincides with our previous publication that the Dpr12/DIP-δ interaction is required for axon regrowth and MB circuit reformation (Bornstein et al., 2021). Moreover, in a parallel ongoing project, we are using proximity labeling to identify binding partners of Dpr12/DIP-δ, and our preliminary results identified CadN as one of the top DIP-δ-binding candidates (manuscript in preparation).

*Drosophila* CadN is an evolutionarily conserved transmembrane protein of the classic cadherin family, with a large and complex extracellular domain. It mediates cell adhesion by homophilic binding, and interacts with the cytoskeleton via its cytoplasmic catenin-binding domain. CadN has been implicated in various neurodevelopmental processes, such as axon guidance, target selection and synaptogenesis (Yonekura et al., 2006; Nern et al., 2005). To genetically examine whether CadN is linked to Dpr4, we used CRISPR/Cas9 to generate a novel CadN mutant allele which lacks its entire coding gene region (*cadN*^ΔORF^). While homozygous mutant clones for *cadN*^ΔORF^ displayed initial growth of both vertical and medial lobes (Fig. 6A-B), they failed to regrow the adult γ-lobe (Fig. 6D vs. C), thereby recapitulating the RNAi-induced phenotype we previously reported (Fahdan et al., 2026). Remarkably, while Dpr4-overexpressing clones consistently showed pruning defects (Fig. 6E), when, on this background, we now mutated CadN within the clones, this was sufficient to suppress the Dpr4 overexpression phenotype, and all axons pruned normally (Fig. 6F). The most plausible interpretation of this finding is that CadN is required to mediate the pruning defect induced by Dpr4 overexpression.

**Figure 6.**
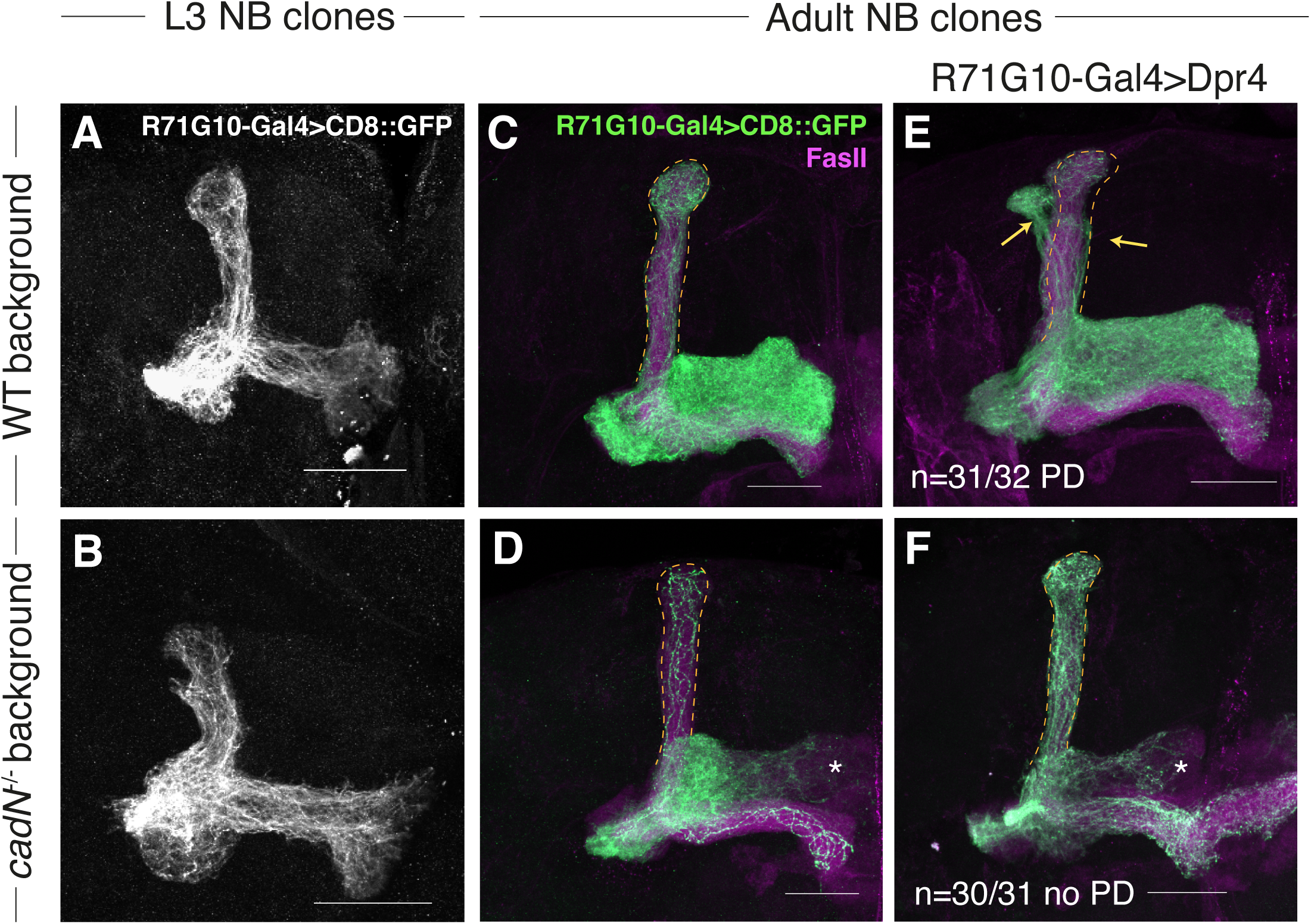
Loss of N-Cadherin suppresses the Dpr4-induced pruning defect. (A-B) Confocal z-projections of 3^rd^ instar larval (L3) MBs with neuroblast (NB) γ-KC MARCM clones, either control (A) or homozygous mutant for the *cadN*^ΔORF^ allele (B). The clones are positively labeled by R71G10-Gal4-driven membranal GFP (CD8::GFP; greyscale). (C-F) Confocal z-projections of adult MBs with neuroblast (NB) γ-KC MARCM clones, either control (C), or overexpressing R71G10-Gal4-driven Dpr4 with (F) or without (E) being homozygous mutant for the *cadN*^ΔORF^ allele, or only homozygous mutant for the *cadN*^ΔORF^ allele (D). The clones are positively labeled by R71G10-Gal4-driven membranal GFP (CD8::GFP; green). Magenta is FasII staining, which strongly labels α/β-axons and weakly labels γ-axons. The α lobe is outlined in orange. Yellow arrows point to unpruned larval axons. Asterisks indicate axon regrowth defects. The fraction of MBs that showed the presented phenotype are indicated in E and F. Scale bar represents 30μm.

## Discussion

One of the least understood aspects of neuronal remodeling is how spatiotemporal precision is achieved. Here, we show that overexpression of the IgSF protein Dpr4 induces a branch-specific axon pruning defect without affecting axon regrowth of the second branch - offering a unique opportunity to explore the spatial regulation of remodeling. Our combined findings, based on genetic interaction and protein engineering experiments, support a model in which overexpressed Dpr4 in MB γ-KCs interacts in *trans*, via its Ig1 domain, with DIP-θ in dopaminergic neurons innervating the vertical lobe. In addition, it likely interacts with another binding partner via its Ig2 domain, to mediate the local inhibition of axon pruning. While mutating the adhesion molecule N-Cadherin (CadN) suppresses the Dpr4-induced pruning defect, it remains to be determined whether the two proteins are part of the same complex (either directly or indirectly interacting), or that CadN acts further downstream of Dpr4. Overall, our study suggests that the spatial specificity of remodeling is likely controlled, at least in part, by cell-cell adhesion, and provides the first genetic evidence of CadN as a potential mediator of Dpr/DIP interactions within a developing neural circuit.

While the observed Dpr4 phenotype results from ectopic overexpression, its unique nature could provide valuable insights into fundamental principles that underlie spatially-restricted neurite elimination, which is prevalent in the developing nervous systems of both vertebrates and invertebrates. The mammalian developing neuromuscular junction is a well-characterized case of non-stereotypic remodeling, in which ongoing competition drives strengthening of a single branch and dismantling of its sister branches. This process is characterized by branch-specific microtubule destabilization mediated, at least in part, by the microtubule-severing enzyme spastin (Brill et al., 2016), however what initiates this branch-specific spastin activation remains unclear. Another classic example is the stereotyped reorganization of corticospinal tract (CST) axons in layer 5 of the mammalian neocortex. Initially neurons from both the motor and visual cortex project axons to identical targets in the spinal cord and the superior colliculus, but during early postnatal life exuberant connectivity is resolved in a branch-specific manner, in which neurons from the motor cortex prune their axonal projections to visual regions and vice versa (O’Leary & Stanfield, 1986). Here, Plexin/Neuropilin signaling was suggested to selectively regulate the pruning of the visual, but not motor, CST axon collaterals, likely via interactions with Semaphorin localized to the targets of visual axons (Low et al., 2008). In *Drosophila*, sensory dendritic arborization (da) neurons that tile the larval body wall are known to completely prune their dendritic arbor during metamorphosis, while their axons remain intact (Yu & Schuldiner, 2014). While it has been shown that the L1-type cell adhesion molecule Neuroglian (Nrg) must be downregulated for pruning to occur (Zhang et al., 2014), its internalization occurs in both dendrites and axons, and thus cannot solely account for the compartment-specific neurite elimination. It is possible that spatial specificity is ensured by intercellular interactions (mediated by Nrg or other CSPs) with adjacent cells, such as epidermal cells or glia, selectively in the dendritic or axonal compartment. Interestingly, larval da neurons were also shown to express many Dprome members (Wang et al., 2022). Whether adhesion molecules provide the link between upstream signals, such as Semaphorin-Plexin signaling, and downstream cytoskeletal rearrangements, is an interesting topic for future investigations.

Shifting back to MB remodeling, one unresolved question regarding its spatial control is why does pruning of γ-axons stop exactly at their branchpoint? Why does it not stop within the lobes or, alternatively, extend distally into the axonal peduncle? The γ-axon branchpoint correlates with a discrete axonal zone, known as γ2, which is innervated by defined PPL1-DANs and MBONs (Aso et al., 2014). Developing γ-KCs endogenously express a wide variety of Dprs (see Fig. 1B), while adult PPL1-DANs innervating the γ2 zone were shown to express several DIPs (most prominently DIP-β; Aso et al., 2019). Thus, our findings that Dpr/DIP-mediated γ-KC-to-PPL1-DAN interactions dictate local inhibition of pruning, could suggest that similar trans-neuronal interactions, involving Dpr/DIPs or similar CSPs, are at play during axon elimination arrest at the branchpoint.

Another intriguing finding in our study is the normal regrowth of the adult medial axonal branch despite persistence of the larval vertical branch. This suggests that pruning and regrowth can be independently controlled as spatially-separable processes at the level of the single branch. As microtubule disassembly of both axonal lobes was shown to be one of the earliest observable steps of the pruning program (Watts et al., 2003), it would be interesting to examine whether Dpr4 overexpression inhibits microtubule disassembly specifically in the vertical lobe. Moreover, how these tempo-chimeric neurons integrate into the adult MB circuitry warrants further investigation. Does the vertical lobe maintain its larval connectivity while the medial lobe forms adult-specific connections? Or do unpruned vertical axons lose their larval connectivity and later ectopically connect with adult-specific neurons within the MB circuit? Notably, during metamorphosis, specific PPL1-DANs innervating the larval vertical γ-lobe remodel their processes to innervate non-MB targets (a process known as trans-differentiation), while newly-born PPL1-DANs innervate the adult vertical α- and α’-lobes (Truman et al., 2023). It remains to be determined whether Dpr4-induced unpruned vertical axons maintain their connectivity to larval PPL1-DANs – thereby preventing their trans-differentiation. Uncovering the functional and behavioral consequences of such chimeric larval/adult circuit connectivity is a fascinating direction for future research.

As IgSF proteins, Dpr/DIPs are considered, at least in principle, to be adhesion-like molecules. We previously showed that adhesion between γ-axons, mediated by the IgSF molecule FasII, must be downregulated for pruning to occur normally (Bornstein et al., 2015). Several studies demonstrated that the Dpr/DIP binding interface depends solely on the Ig1 domain (Cosmanescu et al., 2018; Sergeeva et al., 2020; Carrillo et al., 2015). Therefore, our finding that the Dpr4 overexpression phenotype seems to rely on an Ig2 domain from the Dpr family, suggests that it is not merely Ig1-mediated Dpr::DIP adhesion that physically prevents pruning in a spatially confined manner. As Dpr/DIPs are GPI-anchored and are thus incapable of intracellular signaling, we speculate that a third interacting member - a co-receptor - binds Dpr4, likely via Ig2, to mediate downstream events. This is consistent with recent findings that the Ig2 of the nematode sole DIP, RIG-5, is required for interacting with multiple proteins (Nawrocka et al., 2026). Additionally, the fact that replacing the Ig2 of Dpr4 with that of Dpr12 has the same phenotypic outcome, suggests that this co-receptor(s) is conserved across the Dprome. This is in accordance with our previously published finding, that the Dpr12-DIP-δ interaction can be replaced by Dpr6/10-DIP-α interactions to similarly mediate MB circuit reassembly (Bornstein et al., 2021). With the exception of one recent study that used co-immunoprecipitation to identify the intracellular protein Nocte as Dpr10-associated protein during motor axon pathfinding (Lobb-Rabe et al., 2022), very little is known about the downstream effectors of Dpr/DIP interactions. Our overexpression-based findings provide the first genetic indication of CadN as a potential mediator of Dpr/DIP function. Whether CadN directly interacts with Dprs or DIPs, and if so, whether such potential interaction occurs via Ig2, remains to be determined. While further research is required, we speculate that CadN mediates the local inhibition of axon pruning via its link to catenins and the cytoskeleton. Our findings offer a remarkable starting point to thoroughly explore this potential direct or indirect interaction in more physiological contexts within developing *Drosophila* circuits. Finally, Dpr/DIPs have vertebrate homologs called IgLONs (Cheng et al., 2019), which have been associated with neurodegenerative disorders (Salluzzo et al., 2023), and CadN is conserved in vertebrates (Yonekura et al., 2006). Thus, future research could reveal whether similar interactions contribute to neurodevelopmental processes in higher organisms.

## Materials and methods

### Drosophila melanogaster rearing and strains

All fly strains were reared under standard laboratory conditions at 25 °C on molasses-containing food. Males and females were chosen at random. For developmental analysis, white pupae were collected and incubated for the indicated number of hours before brain dissection. For adult analysis, brains were dissected 3–5 days after eclosion.

UAS-Dpr4.ORF.3xHA was obtained from FlyORF (#F002762). DIP-θ-T2A-Gal4 and UAS-Dpr10D.NV5 were a generous gift from Larry Zipursky, UCLA (Xu et al., 2018). UAS-mtdT.HA was a generous gift from Christopher Potter, Johns Hopkins. R71G10-QF2 and UAS-Dpr12 were previously generated by the Schuldiner lab (Bornstein et al., 2021). R71G10-Gal4 on the 2^nd^ chromosome was previously generated by the Schuldiner lab (Alyagor et al., 2018). The SPARC-L lines (see *Full Drosophila genotypes* below) were previously generated by the Clandinin lab (Currier & Clandinin, 2025). UAS-Dpr4 RNAi (GD13088) was obtained from the Vienna Drosophila Resource Center (VDRC #28518). The following lines were obtained from the Bloomington Drosophila Stock Center (BDSC): R71G10-Gal4 (#39604); TH-Gal4 (#8848); UAS-CD8::GFP (#32186); QUAS-mtdTomato-3xHA (#30004); UAS-DIP-θ RNAi (TRiP JF03069; #28654); UAS-Luciferase VALIUM10 control (#35788); attP40 insertion site control (Msp300; #36304); UAS-EcR-B1^DN^ (UAS-EcR.B1-C655.W650A; #6872); nSyb-IVS-phiC31 on the X (#606050) and 2^nd^ (#84152) chromosomes.

### Design and in vitro testing of Dpr4 mutations that disrupt DIP binding

We analyzed the Dpr::DIP interfaces using either the crystal structure of the Dpr4::DIP-η complex (PDBID: 6EG0; Cosmanescu et al., 2018) or the AF3Complex models (Feldman & Skolnick, 2025) of Dpr4::DIP-θ and Dpr4::DIP-ι. To quantify residue burial, we calculated relative solvent accessibility (RSA=ASA/MaxASA, where ASA is the solvent accessible surface area and MaxASA is the maximum possible solvent accessible surface area for the residue). Ile87 is completely buried in all three complexes (RSA=0%), while Tyr95 and Gln130 are nearly buried (RSA=2-5%); each located in a hydrophobic environment at the Dpr::DIP interface, see Fig.3). Substituting these residues with a negatively charged aspartate was expected to be strongly destabilizing. This design strategy follows established approaches used to abolish binding in other cell-adhesion molecules by introducing buried charge substitutions at conserved hydrophobic interfaces (Lee et al., 2025; Xu et al., 2018; Morano et al., 2025).

To validate our predictions experimentally, we compared the binding of the Dpr4 mutants (I87D, Y95D, Q130D) to that of WT Dpr4 using surface plasmon resonance (SPR; Table 1). All three mutations abolished binding to Dpr4’s cognate partners (DIP-η and DIP-θ) without introducing affinity for a non-cognate partner (DIP-α). The mutation effects listed in Table 1 correspond to ΔΔG > 0.9-1.4 kcal/mol (ΔΔG = RT ln (K_D_(mut)/ K_D_(WT))). Protein purification and SPR were performed as described in previous work (Cosmanescu et al., 2018).

Free energy perturbation (FEP) simulations with replica exchange solute tempering (100 ns sampling, OPLS4 force field, Schrodinger 2022-1 release) further supported these results, predicting substantial loss of affinity: ΔΔG = 6.6 kcal/mol (I87D), 2.6 kcal/mol (Y95D), and 1.6 kcal/mol (Q130D) in the Dpr4::DIP-8 complex. Based on these predictions, we selected the two most destabilizing mutations (I87D and Y95D) for *in vivo* testing.

### Generation of transgenic constructs and transgenic flies

#### Generation of WT and mutant UAS-HA.Dpr4

We inserted the sequence encoding 3 repeats of the HA tag into the WT coding sequence of Dpr4 (Dpr4^WT^, as annotated in FlyBase; https://flybase.org/download/sequence/FBgn0053512/CDS), between the two prolines in positions 34 and 35 (downstream of the predicted signal peptide cleavage site between amino acids 31-32, and upstream of Ig1). The HA-Dpr4 sequence was cloned into the Gateway entry vector pTwist ENTR (Twist Bioscience), and then recombined into the pDEST-UAS-IVS-Syn21-p10 destination vector (Rabinovich et al., 2016) using LR recombinase (Invitrogen).

UAS transgenes harboring mutations that disrupt DIP binding, or that harbor Ig2 replacements, were designed and cloned in the same manner, except introducing the following changes:

1. UAS-HA.Dpr4^I87D^: Isoleucine in position 87 (ATC) replaced with Aspartate (GAC).
2. UAS-HA.Dpr4^Y95D^: Tyrosine in position 95 (TAC) replaced with Aspartate (GAC).
3. UAS-HA.Dpr4^Ig2-Dpr12^: The Ig2 domain was defined as the entire sequence residing between the two cysteines in positions 170 and 227, and was replaced with the Ig2 domain of Dpr12 isoform C (as annotated in FlyBase) - also encompassing the entire sequence between the two conserved cysteines of Ig2 (in positions 208 and 267).
4. UAS-HA.Dpr4^Ig2-FasII^: The Ig2 domain of Dpr4 (between cysteines 170 and 227) was replaced with the sequence encoding the Ig2 domain of FasII isoform C, spanning between the cysteines in positions 159 and 207.

Notably, following the destruction of our lab (on June 15^th^, 2025) the original UAS-HA.Dpr4^WT^ line was lost, and we therefore regenerated it using the services of VectorBuilder, by cloning the WT HA-Dpr4 sequence into their *Drosophila* ΦC31-based vector pUASTattB. These two lines display the exact same phenotype and are used interchangeably throughout the manuscript (often referred to as UAS-HA.Dpr4^WT^).

All UAS-HA.Dpr4 plasmids (WT and mutant) were injected to *Drosophila* embryos and integrated into the attP40 landing site using ΦC31 integration (Bestgene).

#### Generation of QUAS-Dpr4

In parallel to the generation of UAS-HA.Dpr4, an untagged UAS-Dpr4 plasmid and transgenic fly was generated in a similar manner (i.e., Dpr4 CDS in pDEST-UAS-IVS-Syn21-p10; does not appear in the manuscript). This plasmid provided the template for the generation of the QUAS-Dpr4 plasmid, using restriction free (RF) cloning. A first PCR reaction was performed to generate a megaprimer, with UAS-Dpr4 providing the template, using the following primers:

Forward: GGGAATTCGTTAACAGATCTGCGGCCGCGCCAGGCTATGTGGACAACGGAC

Reverse: CTCTGTAGGTAGTTTGTCCAATTATGTCACATTTTTCGTAATCTTCCTAATG

The megaprimer was then used in a second PCR reaction to insert the Dpr4 CDS into the pQUAST-attB target plasmid - a generous gift from Christopher Potter, Johns Hopkins.

The QUAS-Dpr4 plasmid was injected to *Drosophila* embryos and integrated into the attP40 landing site using ΦC31 integration (Bestgene).

Important to note that a 3xHA-tag sequence was added at the C-terminal end of Dpr4, to mimic the original UAS FlyORF transgene (and since we generated the QUAS-Dpr4 plasmid prior to knowing that Dpr4 is GPI-anchored). Based on our results (SFig. 2), this HA tag is not predicted to be a part of the final protein. Following the destruction of our lab we generated a new, N-terminal tagged QUAS-HA-Dpr4 transgene, following a similar strategy of the new UAS-HA-Dpr4^WT^ (using the services of VectorBuilder, with their *Drosophila* ΦC31-based vector pUASTattB as the backbone, modified to replace the UAS promoter with a 5XQUAS promoter). This transgenic fly recapitulates the phenotype observed using the original QUAS-Dpr4 fly line that no longer exists, but also shows HA labeling throughout the entire neuron. However, the data currently presented in Figure 4 is with the original QUAS-Dpr4 fly.

#### Generation of the CadN^ΔORF^ allele

To delete the entire coding gene region of CadN, we used the FlyCRISPR algorithm (https://flycrispr.org/) to select the following gRNAs:

N-ter: AAAAGGTGTTAATACCAAAC

C-ter: CAATACCGAACTAGAATTGT

Both gRNAs were cloned into the pCFD4 plasmid (Addgene #49411) using transfer-PCR (TPCR; Meltzer et al., 2019; Erijman et al., 2014). The CadN-pCFD4 plasmid was injected into the attP40 landing site using ΦC31 integration (BestGene). Injected flies were crossed with nanos-Cas9 flies (BDSC #54591). After two generations, single males were crossed with balancers and checked for deletion using the following primers:

Forward: GGAGAAGGTAGACTGGACAAAGG

Reverse: TACCGGTTGCCCTAACAACTAAA

The resulting *CadN^ΔORF^*allele is a deletion of 89,143bp - leaving only 54bp of the gene’s coding sequence: 31bp upstream + 23bp downstream of the deletion (including the ATG and stop codon).

#### Generation of MARCM clones

MARCM clones of γ-KCs were generated by a 30-60 minute heat-shock at 37°C of newly hatched larvae, 24 hours after egg laying, as previously described (Lee & Luo, 1999).

#### Immunostaining and imaging

*Drosophila* brains were dissected in cold Ringer’s solution, fixed using 4% paraformaldehyde (PFA) for 20 minutes in room temperature (RT), and washed in phosphate buffer with 0.3% Triton-X (PBT; three quick washes followed by three 20-minute washes). Non-specific staining was blocked using 5% normal goat serum (NGS) in PBT for 30 minutes in RT, and brains were then subjected to primary antibody staining overnight at 4°C. Primary antibodies included chicken anti-GFP 1:500 (GFP-1020; AVES), mouse anti-FasII 1:25 (1D4; DSHB), rat anti-HA 1:100 (11867423001; Sigma Aldrich), rabbit anti-TH 1:500 (AB152; Merck Millipore), and rat anti-RFP 1:1000 (5F8; ChromoTek). Brains were rinsed (3 quick washes and 3 × 20-minute washes) and stained with secondary antibodies for 2 hours at RT. Secondary antibodies included FITC donkey anti-chicken 1:300 (703-095-155; Jackson immunoresearch), Alexa fluor 647 goat anti-mouse 1:300 (A-21236; Invitrogen), Alexa fluor 647 goat anti-rabbit 1:300 (A-21236; Invitrogen), Alexa fluor 647 goat anti-rat 1:300 (A-21247; Invitrogen), and Alexa fluor 568 goat anti-rat 1:300 (A-11077; Invitrogen).

For immunostaining of non-permeabilized brains, brains were dissected in ice-cold Ringer’s solution (including removal of all trachea), and then incubated for 3h in 4°C with the primary antibody (rat anti-HA 1:100) diluted in Ringer’s solution. Following incubation, brains underwent 3 × 15-minute washes with Ringer’s solution in 4°C, and then fixed in 4% PFA in Phosphate-Buffered Saline (PBS) for 20 minutes at RT, followed by 3 × 20 minute washes with PBS in RT. Brains were then blocked with 5% NGS in PBS for 30 minutes in RT, incubated with the secondary antibody (Alexa fluor 647 goat anti-rat 1:300) diluted in PBS for 3h in RT, and washed 3 times with PBS before mounting.

All brains were mounted on Slowfade (S-36936; Invitrogen) and imaged with Zeiss LSM 980 or Zeiss LSM 990 confocal microscopes. Images were processed with ImageJ (NIH).

#### Quantification and statistical analysis

The severity of pruning defects in the MBs of adult flies (Fig. 1J, Fig. 3F, Fig. 4G, Fig. 5D) was blindly ranked by an independent investigator, on a scale of 0 (WT-like phenotype) to 4 (severe pruning defect), as routinely done in the Schuldiner lab (e.g., Marmor-Kollet et al., 2023; Mayseless et al., 2023; Keret et al., 2025). Groups were compared by Kruskal-Wallis test followed by Mann-Whitney U-test with FDR correction for multiple comparisons.

Quantification of branch-specific pruning (Fig. 2D, SFig. 4D) was performed using FIJI, by measuring the mean GFP intensity within a selected ROI of 50×50 pixels, in a sub-z-stack spanning the entire lobe in the Z-axis (normally in the range of 10-30 slices) - in both the vertical and the medial lobes as annotated in SFig. 4A-C (yellow and magenta boxes, respectively). The mean intensity values (listed after plotting the Z-axis profile) were averaged across all slices within a given ROI, and the vertical/medial mean intensity ratio was calculated based on these average values for each image individually. The three groups were compared using Welch’s Anova, and Games Howell post hoc test for pairwise comparisons.

Statistical analyses were conducted in R version 4.4.3 (R Core Team, 2025). Specific p-values are reported within the relevant figure legends.

#### Full Drosophila genotypes

hsFLP is y,w,hsFLP122; mtdT is myr::tdTomato-3xHA; 40A and G13 are FRTs on 2L and 2R respectively; G80 is TubP-Gal80. attP40 control is the empty attP40 landing site (BDSC #36304); UAS-EcR^DN^ is UAS-EcR.B1-C655.W650A (BDSC #6872); UAS-Dpr4.ORF is UAS-Dpr4.ORF.3xHA. Abbreviations of the UAS-HA.Dpr4 transgenes are explained in the Materials and Methods section *Generation of WT and mutant UAS-HA.Dpr4*. Males and females were used interchangeably but only the female genotype is written.

<u>Figure 1:</u>

1D: y,w/y,v; UAS-CD8::GFP/attP40 control; R71G10-Gal4/+

1E: y,w; UAS-CD8::GFP/UAS-HA.Dpr4^WT^; R71G10-Gal4/+

1F+H: hsFLP, UAS-CD8::GFP/+; R71G10-Gal4,G13,G80/40A,G13,cn,bw

1G+I: hsFLP, UAS-CD8::GFP/+; R71G10-Gal4,G13,G80/40A,G13,cn,bw; +/UAS-Dpr4.ORF

<u>Figure 2:</u>

2A: y,w/y,v; UAS-CD8::GFP/attP40 control; R71G10-Gal4/+

2B: y,w/w; UAS-CD8::GFP/UAS-EcR-B1^DN^; R71G10-Gal4/+

2C: y,w; UAS-CD8::GFP/UAS-HA.Dpr4^WT^; R71G10-Gal4/+

2D: nSyb-IVS-phiC31/20xUAS-SPARC2-S-LexA; R71G10-Gal4/13xLexAop-SPARC2-I-GCaMP8-P2A-mtdT

2E: nSyb-IVS-phiC31/w; UAS-SPARC2-S-lexA/UAS-EcR-B1^D^; R71G10-Gal4/LexAop-SPARC2-S-GCaMP8m

2F: nSyb-IVS-phiC31/w; UAS-SPARC2-S-lexA/UAS-HA.Dpr4^WT^; R71G10-Gal4/LexAop-SPARC2-S-GCaMP8m

<u>Figure 3:</u>

3B: y,w/y,v; UAS-CD8::GFP/attP40 control; R71G10-Gal4/+

3C: y,w; UAS-CD8::GFP/UAS-HA.Dpr4^WT^; R71G10-Gal4/+

3D: y,w; UAS-CD8::GFP/UAS-HA.Dpr4^I87D^; R71G10-Gal4/+

3E: y,w; UAS-CD8::GFP/UAS-HA.Dpr4^Y95D^; R71G10-Gal4/+

<u>Figure 4</u>:

4B: y,w/y,v; R71G10-QF2,QUAS-mtdT/attP40 control

4C: y,w; R71G10-QF2,QUAS-mtdT/QUAS-Dpr4

4D: y,w or y,v/y,w or w; R71G10-QF2,QUAS-mtdT/QUAS-Dpr4; UAS-DIP-θ RNAi/TH-Gal4

4E: y,w or y,v/y,w or w; R71G10-QF2,QUAS-mtdT/QUAS-Dpr4; UAS-Luciferase/TH-Gal4

4F: y,w or y,v/y,w; R71G10-QF2,QUAS-mtdT/+; UAS-Luciferase/TH-Gal4

<u>Figure 5:</u>

5A: y,w; UAS-CD8::GFP/UAS-HA.Dpr4^WT^; R71G10-Gal4/+

5B: y,w; UAS-CD8::GFP/UAS-HA.Dpr4^Ig2-Dpr12^; R71G10-Gal4/+

5C: y,w; UAS-CD8::GFP/UAS-HA.Dpr4^Ig2-FasII^; R71G10-Gal4/+

<u>Figure 6:</u>

6A,C: hsFLP, UAS-CD8::GFP/+; G80,40A/40A,G13,cn,bw; R71G10-Gal4/+

6B,D: hsFLP, UAS-CD8::GFP/+; G80,40A/*ncad*^ΔORF^,40A,G13,cn,bw; R71G10-Gal4/+

6E: hsFLP, UAS-CD8::GFP/+; G80,40A/40A,G13,cn,bw; R71G10-Gal4/UAS-Dpr4.ORF

6F: hsFLP, UAS-CD8::GFP/+; G80,40A/*ncad*^ΔORF^,40A,G13,cn,bw; R71G10-Gal4/UAS-Dpr4.ORF

<u>SFigure 1:</u>

S1A: y,w/w; UAS-CD8::GFP/+; R71G10-Gal4/+

S1B: y,w/w; UAS-CD8::GFP/UAS-Dpr4 RNAi (GD13088); R71G10-Gal4/+

<u>SFigure 2:</u>

S2A: y,w/y,v; UAS-CD8::GFP/attP40 control; R71G10-Gal4/+

S2B: y,w/+; UAS-CD8::GFP/+; R71G10-Gal4/UAS-Dpr4.ORF

S2C+D: y,w; UAS-CD8::GFP/UAS-HA.Dpr4^WT^; R71G10-Gal4/+

<u>Figure 3:</u>

S3A: y,w/y,v; UAS-CD8::GFP/attP40 control; R71G10-Gal4/+

S3B: y,w; UAS-CD8::GFP/UAS-Dpr10D.NV5; R71G10-Gal4/+

S3C: y,w; UAS-CD8::GFP/+; R71G10-Gal4/UAS-Dpr12

<u>SFigure 4:</u>

S4A: y,w/y,v; UAS-CD8::GFP/attP40 control; R71G10-Gal4/+

S4B: y,w/w; UAS-CD8::GFP/UAS-EcR-B1^DN^; R71G10-Gal4/+

S4C: y,w; UAS-CD8::GFP/UAS-HA.Dpr4^WT^; R71G10-Gal4/+

<u>SFigure 6:</u>

S6A-C: R71G10-QF2,QUAS-mtdT,UAS-CD8::GFP/DIP-θ-T2A-Gal4

<u>SFigure 7:</u>

7A: y,w; UAS-CD8::GFP/UAS-HA.Dpr4^WT^; R71G10-Gal4/+

7B: y,w; UAS-CD8::GFP/UAS-HA.Dpr4^Ig2-Dpr12^; R71G10-Gal4/+

7C: y,w; UAS-CD8::GFP/UAS-HA.Dpr4^Ig2-FasII^; R71G10-Gal4/+

7D+E: y,w; UAS-CD8::GFP/UAS-mtdT; R71G10-Gal4/+

## Acknowledgements

We thank the Bloomington Drosophila Stock Center and the Vienna Drosophila Resource Center for providing fly stocks. Monoclonal antibodies were obtained from the Developmental Studies Hybridoma Bank, created under the auspices of the NICHD and maintained by the University of Iowa. We are grateful to Ron Rotkopf for assistance with statistical analysis; Shifra Ben-Dor for bioinformatics; Larry Zipursky, Kai Zinn and Christopher Potter for kindly sharing valuable reagents; Neta Hanuka from the Schuldiner lab for help with quantifications.

This work was supported by National Science Foundation (NSF) IOS-2321481 (B.H.); NSF-BSF 2023611 (O.S.); and European Research Council (ERC) Advanced Grant 101054886 “NeuRemodelBehavior” (O.S.). O.S. is the incumbent of the Prof. Erwin Netter Professorial Chair of Cell Biology.

