## Supplementary data and figures for "Branch-specific axon pruning induced by Dpr4/DIP-θ transneuronal interactions"

Supplemental Figures

Hagar Meltzer[1*], Shirel Shahar[1*], Alina P. Sergeeva[2,3], Bavat Bornstein[1], Gal Shapira[1], Phinikoula S. Katsamba[4], Seetha Mannepalli[4], Fabiana Bahna[4], Noa Moreno[1], Idan Alyagor[1], Voctoria Berkun[1], Timothy Currier[6], Lawrence Shapiro[2,4], Barry Honig[2,3,4,5], Oren Schuldiner[1#]

[1] Dept. of Molecular Cell Biology and Dept. of Molecular Neuroscience, Weizmann Institute of Science, Rehovot, Israel

[2] Department of Biochemistry and Molecular Biophysics, Columbia University, New York, U.S.A

[3] Department of Systems Biology, Columbia University, New York, U.S.A

[4] Zuckerman Mind Brain Behavior Institute, Columbia University, New York, U.S.A

[5] Department of Medicine, Columbia University, New York, U.S.A

[6] Department of Neurobiology, Stanford University School of Medicine, Stanford, CA, U.S.A

* These authors contributed equally

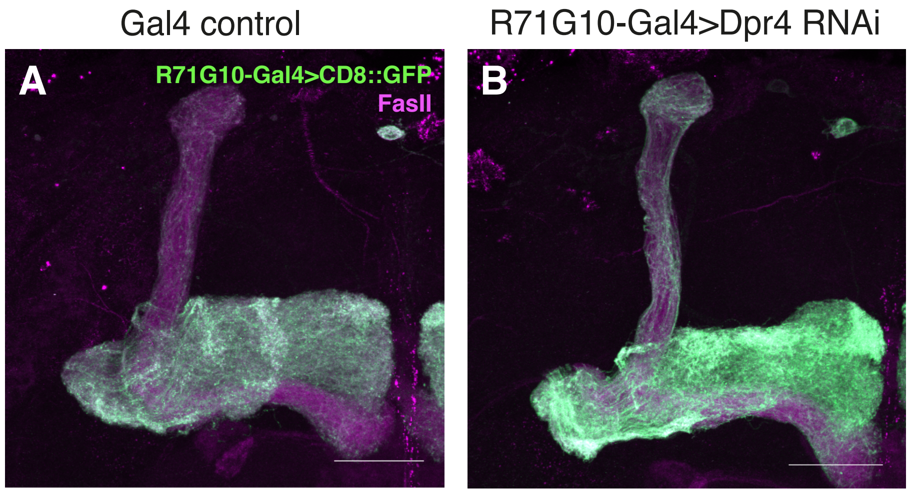

Gal4 control
R71G10-Gal4>Dpr4 RNAi
A
R71G10-Gal4>CD8::GFP
FasII
B

**Supplemental Figure 1. Knocking down Dpr4 does not affect axon pruning or regrowth**

(A-B) Confocal z-projections of adult control MBs in which the largely γ-specific R71G10-Gal4 (also stochastically expressed in α/β-KCs) drives expression of membranal GFP (CD8::GFP; green; A is control), or additionally RNAi targeting Dpr4 (GD13088; B). Magenta is FasII staining, which strongly labels α/β-axons and weakly labels γ-axons. Scale bar represents 30μm.

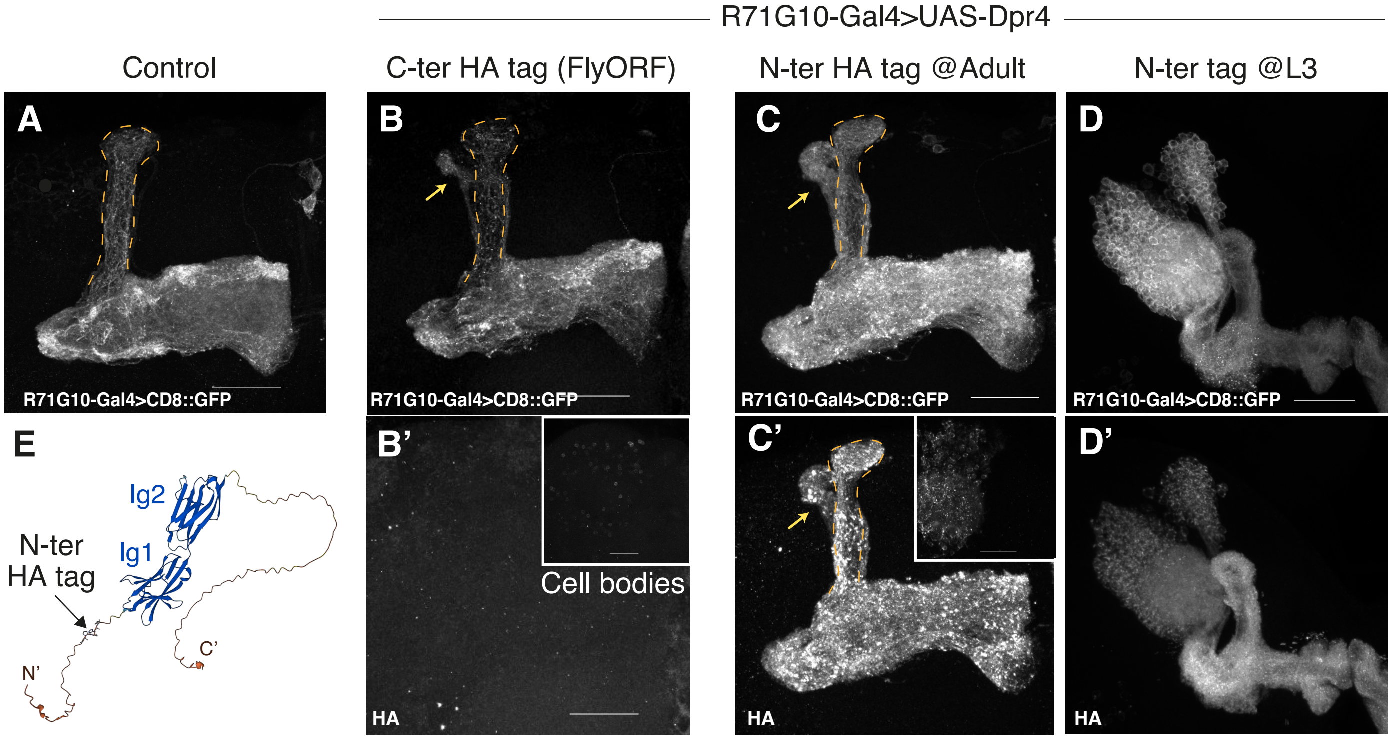

R71G10-Gal4>UAS-Dpr4
Control
C-ter HA tag (FlyORF)
N-ter HA tag @Adult
N-ter tag @L3
A
B
C
D
R71G10-Gal4>CD8::GFP
R71G10-Gal4>CD8::GFP
R71G10-Gal4>CD8::GFP
R71G10-Gal4>CD8::GFP
E
B'
C'
D'
Ig2
Ig1
N-ter
HA tag
N'
C'
Cell bodies
HA
HA
HA

**Supplemental Figure 2. Overexpressed Dpr4 is localized throughout γ-KCs and inhibits their axon pruning**

(A-D) Confocal z-projections of adult (A-C) or 3$^{rd}$ instar larva (L3; D) MBs, in which R71G10-Gal4 drives expression of membranal GFP (CD8::GFP; greyscale in A-D), as well as overexpression of Dpr4 (from the FlyORF collection) with a C-terminal HA tag (B), or a newly-generated Dpr4 transgene with an N-terminal (post signal peptide) HA tag (C-D, schematized in E). The HA signal is absent from axons in the C-ter tagged transgene (B') and only observed in a fraction of the cell bodies (inset in B'), but is widely localized throughout the axons and cell bodies in the new N-ter tagged transgene in both adult (C') and L3 (D'). The α lobe is outlined in orange. Yellow arrows point to unpruned larval axons. Scale bar represents 30μm.

(E) Scheme of the location of the N-terminal HA tag (shown in C'-D'), downstream of the signal peptide and upstream of the 1$^{st}$ immunoglobulin domain (Ig1), between two Prolines. Image of Dpr structure is by AlphaFold (https://alphafold.ebi.ac.uk/entry/Q59DX6).

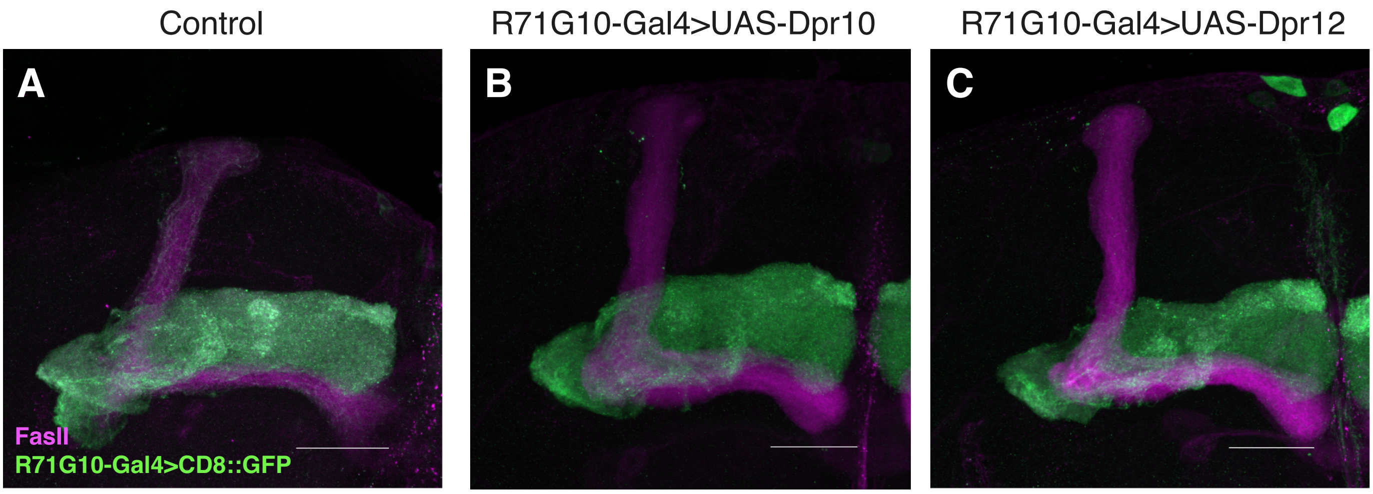

Control
R71G10-Gal4>UAS-Dpr10
R71G10-Gal4>UAS-Dpr12
A
B
C
FasII
R71G10-Gal4>CD8::GFP

**Supplemental Figure 3. Overexpression of other Dprs does not affect pruning**

(A-C) Confocal z-projections of adult MBs in which the γ-specific R71G10-Gal4 drives expression of membranal GFP (CD8::GFP; green), either control (A) or also driving overexpression of Dpr10 (B) or Dpr12 (C). Magenta is FasII staining, which strongly labels α/β-axons and weakly labels γ-axons. Scale bar represents 30μm.

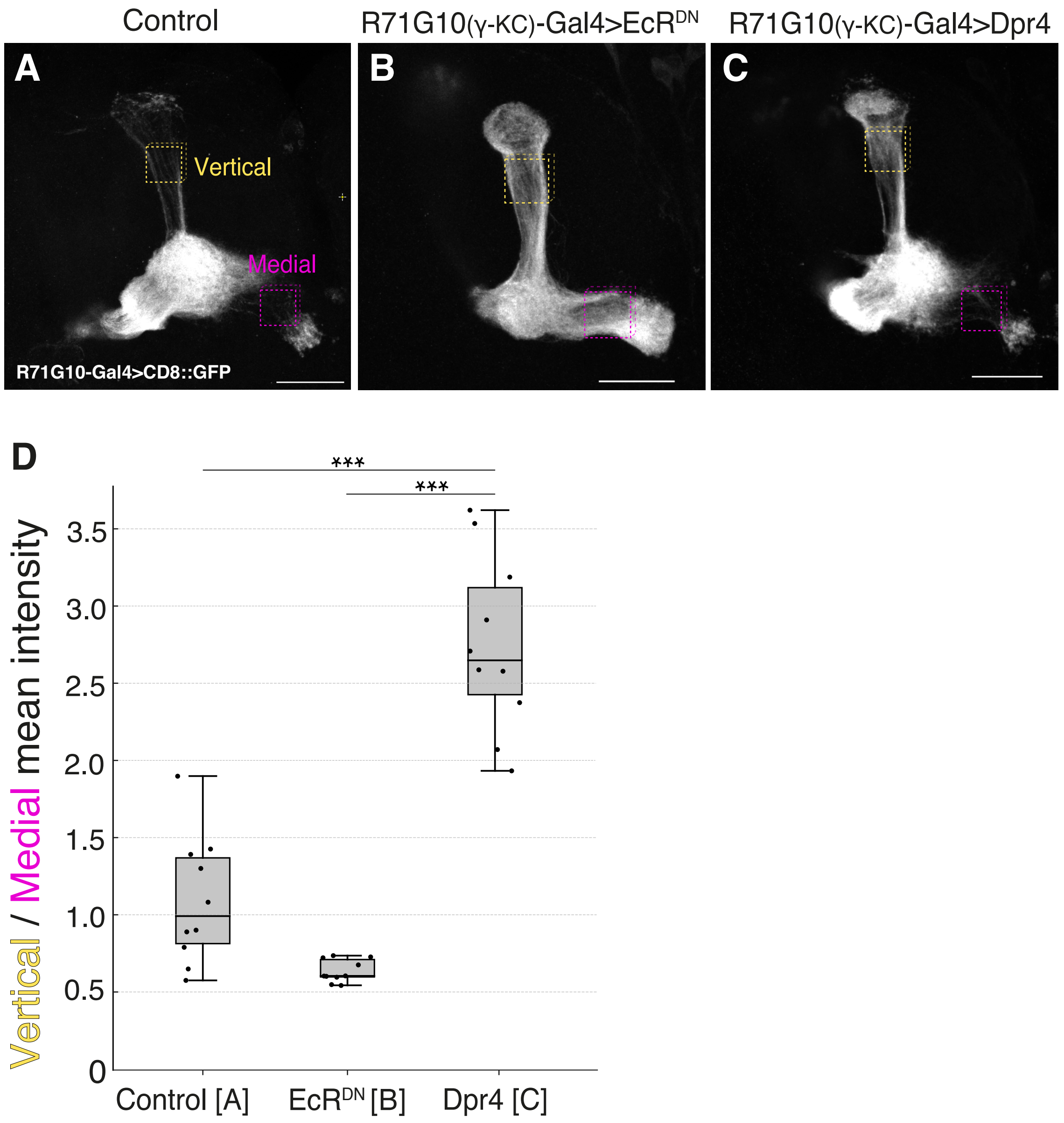
Control
R71G10(γ-KC)-Gal4>EcR^DN
R71G10(γ-KC)-Gal4>Dpr4
A
B
C
Vertical
Medial
R71G10-Gal4>CD8::GFP
D
***
***
Vertical / Medial mean intensity
3.5
3.0
2.5
2.0
1.5
1.0
0.5
0
Control [A]
EcR^DN [B]
Dpr4 [C]

**Supplemental Figure 4. Quantification of branch-specific axon pruning**

(A-C) Confocal z-projections of MBs at 24 hours after puparium formation (h APF), in which the γ-specific R71G10-Gal4 drives expression of membranal GFP (CD8::GFP; greyscale), either in control (A) or also driving overexpression of EcR-B1$^{DN}$ (B) or Dpr4 (C). Yellow arrows point to unpruned γ lobes. The yellow and magenta boxes represent the regions (along the z-stack) taken in the vertical and medial lobes, respectively, for measurements of mean GFP intensity (see quantification in D). Scale bar represents 30μm.

(D) Quantification of the vertical/medial ratio of mean GFP intensities, as measured within the vertical and medial lobes in control, EcR-B1$^{DN}$ and Dpr4 overexpressing γ-KCs (see yellow and pink boxes in A-C as an example, full details in the Materials and Methods section). In each box plot, the horizontal line represents the median, the box spans the interquartile range (IQR; from the 25th to 75th percentiles), and the whiskers extend to the smallest and largest values within 1.5×IQR from the lower and upper quartiles, respectively. Black dots represent the individual data points.

Welch's Anova $p = 1.659 \times 10^{-7}$; Games Howell test: [A] vs. [C] ***$p = 3.28 \times 10^{-6}$; [B] vs. [C] ***$p = 2.01 \times 10^{-6}$

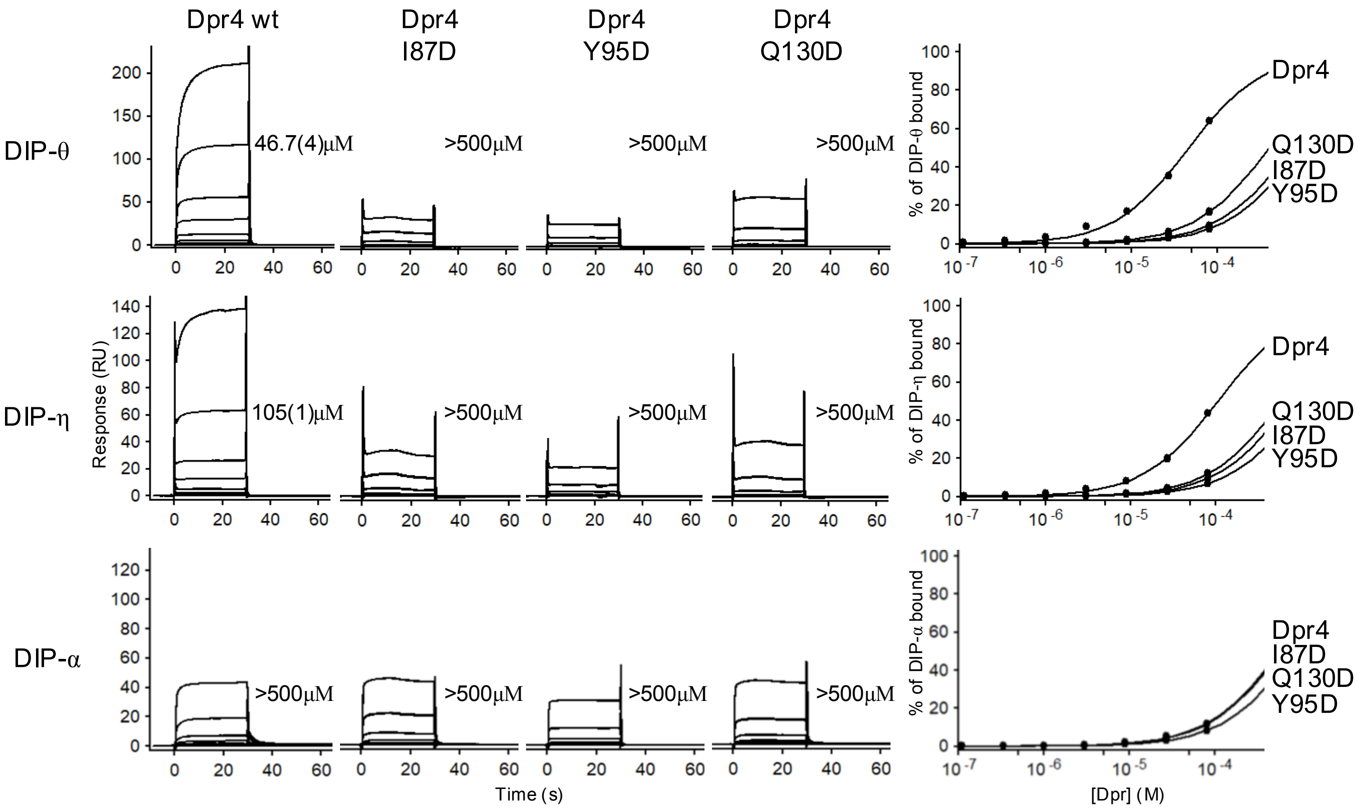

Dpr4 wt
Dpr4 I87D
Dpr4 Y95D
Dpr4 Q130D
DIP-θ
46.7(4)μM
>500μM
>500μM
>500μM
DIP-η
105(1)μM
>500μM
>500μM
>500μM
DIP-α
>500μM
>500μM
>500μM
>500μM
Response (RU)
Time (s)
% of DIP-θ bound
% of DIP-η bound
% of DIP-α bound
[Dpr] (M)
Dpr4
Q130D
I87D
Y95D
Dpr4
I87D
Q130D
Y95D

**Supplemental Figure 5. SPR sensorgrams and corresponding binding isotherms for Dpr4 wild-type (wt) and three Dpr4 mutants used as analytes over immobilized DIP-θ, DIP-η, and DIP-α surfaces**.

Equilibrium dissociation constants ($K_D$) for interactions with cognate (DIP-η and DIP-θ) and non-cognate (DIP-α) DIPs are provided in Table 1. Responses are plotted against the Dpr concentration tested in each experiment. The Dpr concentration corresponding to 50% bound, represents the $K_D$.

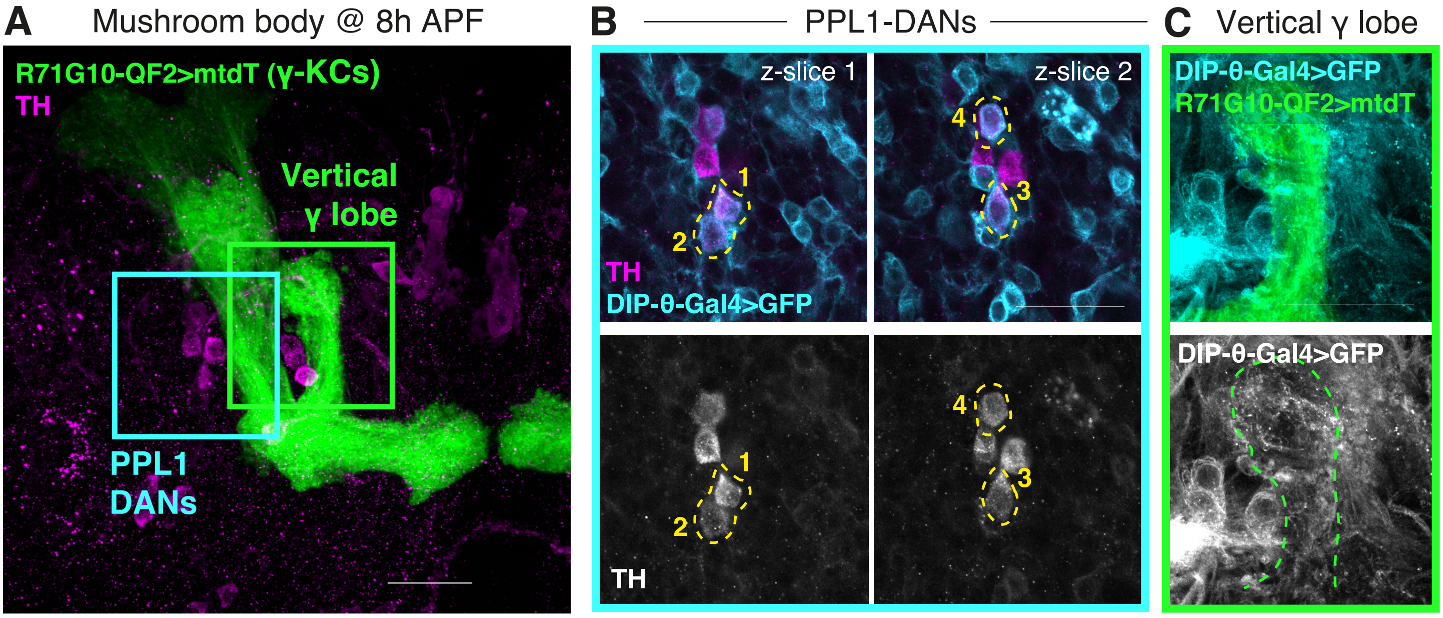
A
Mushroom body @ 8h APF
R71G10-QF2>mtdT (γ-KCs)
TH
Vertical
γ lobe
PPL1
DANs
B
PPL1-DANs
z-slice 1
z-slice 2
1
2
3
4
TH
DIP-θ-Gal4>GFP
1
2
3
4
TH
C
Vertical γ lobe
DIP-θ-Gal4>GFP
R71G10-QF2>mtdT
DIP-θ-Gal4>GFP

**Supplemental Figure 6. DIP-θ is expressed in DANs innervating the vertical lobe at the onset of pruning**

(A) Confocal z-projection of a MB at 8 hours after puparium formation (h APF), in which R71G10-QF2 drives expression of myr::tdTomato (mtdT; green) in γ-KCs, and DIP-θ-T2A-Gal4 drives expression of membranal GFP (CD8::GFP, channel not shown in this panel) in DIP-θ-expressing cells. Magenta is staining for anti-tyrosine hydroxylase (TH, a marker of dopaminergic neurons).

(B) Confocal single slices which are a magnified view of the PPL1-DAN cell body region (cyan box in A). Cyan is DIP-θ-T2A-Gal4-Driven CD8::GFP. Magenta/greyscale is anti-TH staining. Cell bodies that are labelled by both cyan and magenta are outlined in yellow, two different z-slices are displayed to capture all 4 such cells.

(C) Confocal z-projection which is a magnified view of the vertical lobe region (green box in A). Green is R71G10-QF2-driven mtdT. Cyan/greyscale is DIP-θ-T2A-Gal4-Driven CD8::GFP. The vertical γ lobe is outlined in green.

Scale bar represents 30μm.

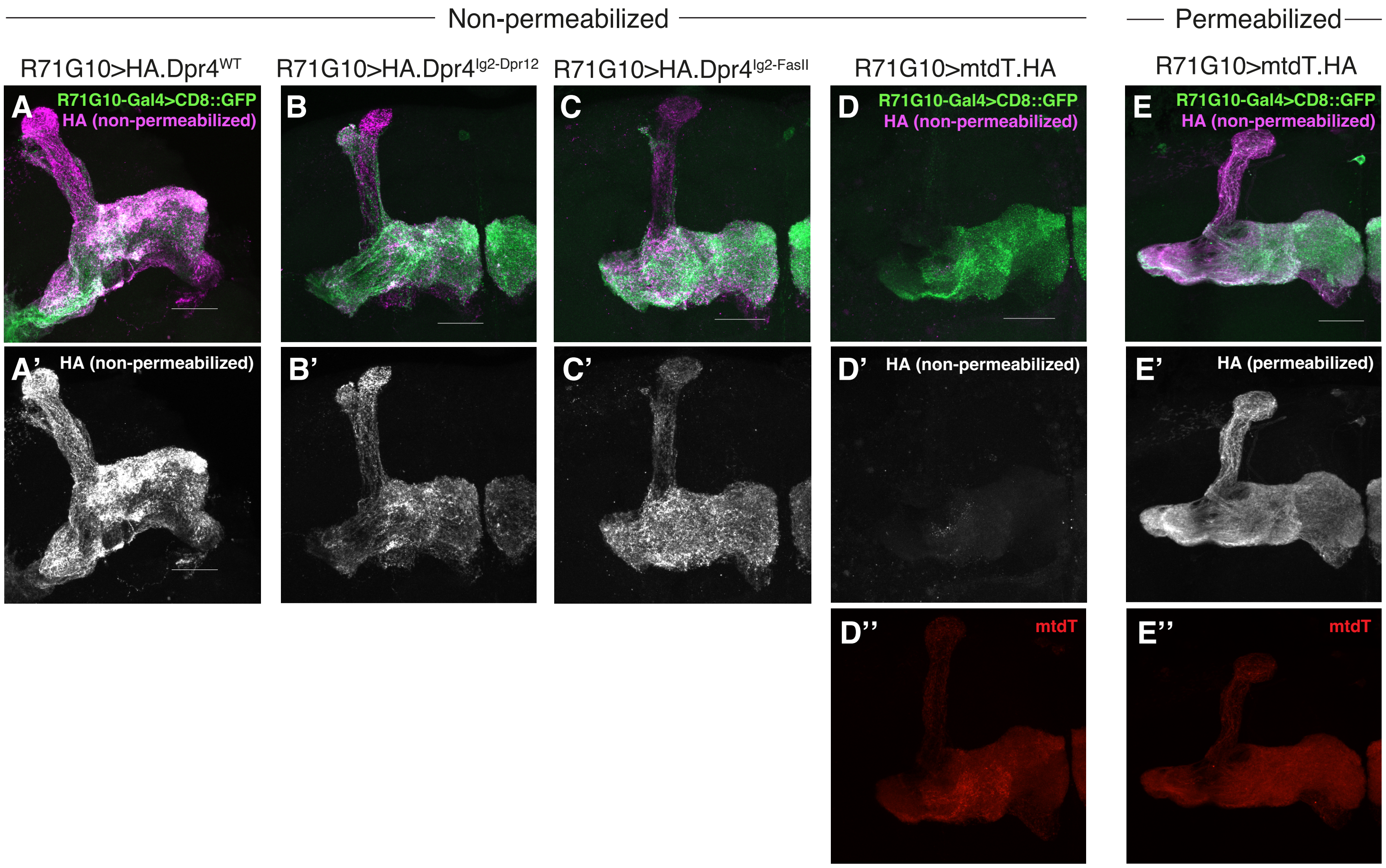
Non-permeabilized
Permeabilized
R71G10>HA.Dpr4WT
R71G10>HA.Dpr4Ig2-Dpr12
R71G10>HA.Dpr4Ig2-FasII
R71G10>mtdT.HA
R71G10>mtdT.HA
A
R71G10-Gal4>CD8::GFP
HA (non-permeabilized)
B
C
D
R71G10-Gal4>CD8::GFP
HA (non-permeabilized)
E
R71G10-Gal4>CD8::GFP
HA (non-permeabilized)
A'
HA (non-permeabilized)
B'
C'
D'
HA (non-permeabilized)
E'
HA (permeabilized)
D''
mtdT
E''
mtdT

**Supplemental Figure 7. Chimeric Ig2-replaced Dpr4 proteins are expressed on the cell surface of γ-KCs**

(A-C) Confocal z-projections of adult MBs in which the γ-specific R71G10-Gal4 drives expression of membranal GFP (CD8::GFP; green) as well as Dpr4, either WT (A), or in which the entire Ig2 is replaced with the Ig2 of Dpr12 isoform C (B) or of FasII isoform C (C). Non-permeabilized staining for the HA tag inserted in the N-terminus of the WT/mutant Dpr4 protein is shown in magenta in A-C or in greyscale in A'-C'.

(D-E) Confocal z-projections of adult MBs in which R71G10-Gal4 drives expression of membranal GFP (CD8::GFP; green) as well as HA-tagged myristoylated tdTomato (mtdT::HA; red in D'' and E''). Staining for the HA tag inserted in the C-terminus of mtdT - which is intracellular and thus used as control - in either non-permeabilized (D) or permeabilized (E) brains is shown in magenta in D-E and in greyscale in D'-E'.

Scale bar represents 30μm.
